# Sialylated Gangliosides are Required for Plasma Membrane Organization and Neuronal Function

**DOI:** 10.64898/2026.03.18.712603

**Authors:** Henry G. Barrow, Zhuang Z. Han, Alex S. Nicholson, Sinéad Strasser, Daniel A. Nash, John O. Suberu, Robin Antrobus, Danielle te Vruchte, David A. Priestman, Stephen C. Graham, Frances M. Platt, Janet E. Deane

## Abstract

Gangliosides are abundant neuronal glycosphingolipids, yet their roles in organizing the plasma membrane and supporting neuronal function remain poorly defined. Mutations in the biosynthetic enzymes ST3GAL5 or B4GALNT1 cause severe neurodevelopmental disorders, yet their cellular consequences are unclear. Using isogenic human iPSC-derived cortical neurons, we show that loss of these enzymes eliminates major neuronal gangliosides but produces strikingly divergent outcomes. ST3GAL5 deficiency reprograms the glycosphingolipid repertoire toward non-neuronal species and abolishes network-level electrical activity. In contrast, B4GALNT1-deficient neurons retain near-normal excitability, supported by accumulation of simple sialylated precursors (GM3/GD3). Quantitative proteomics reveals a profound loss of plasma membrane proteins, including ion channels and synaptic organizers, only in ST3GAL5-deficient neurons. These findings identify sialylated glycosphingolipids as essential scaffolds for plasma membrane organization and neuronal excitability, providing a mechanistic basis for the severe phenotype caused by loss of GM3 synthase in humans.

**TEASER:** Human neuronal models reveal why loss of GM3 synthase causes severe neurodevelopmental disease

## INTRODUCTION

Glycosphingolipids (GSLs) are essential bioactive lipids that are enriched on the cell surface where they play crucial roles in cell recognition, cell identity and membrane protein function[1–3]. GSLs are characterized by structurally diverse glycan headgroups (**Fig.1A**) and interact with cholesterol to form highly ordered membrane microdomains (also known as lipid rafts). These microdomains support the spatial organization and assembly of signaling and adhesion complexes. Although GSLs are also known to influence membrane protein trafficking, the details of how they regulate this process remain unclear[4,5].

**Figure 1:**
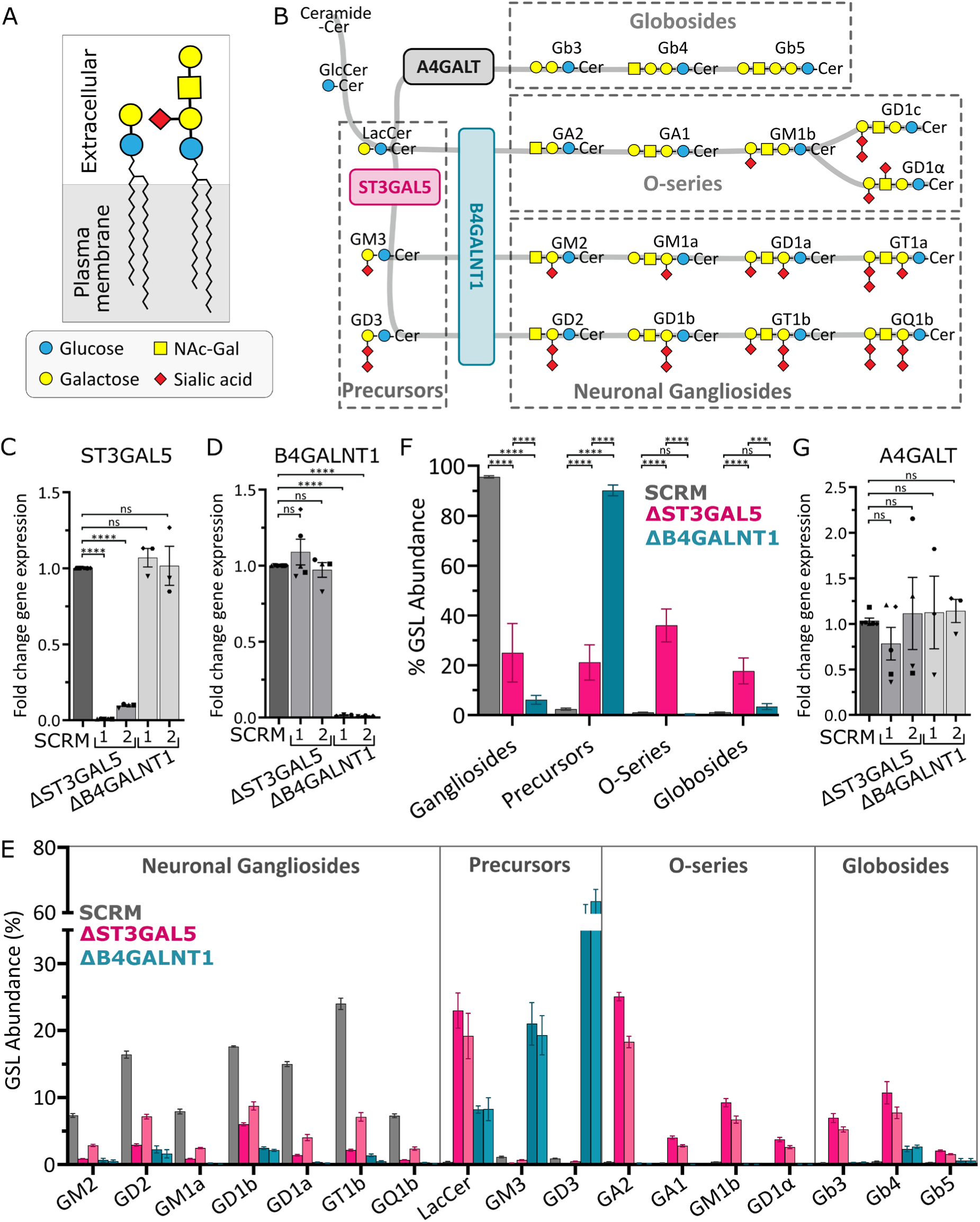
ΔST3GAL5 and ΔB4GALNT1 i3 neurons have severe and contrasting changes to their GSL composition. (A) Simplified diagram of glycosphingolipid (GSL) structure with a membrane embedded ceramide tail and extracellular glycan head groups. (B) Schematic of GSL synthesis with lipid series (dashed boxes) and select enzymes (solid boxes) labelled. qPCR of (C) ST3GAL5, (D) B4GALNT1 and (G) A4GALT gene expression in i3N at 28 dpi. Fold change relative to SCRM controls are shown for each cell line. The mean ± SEM is shown for N = 3-6 biological replicates (squares, triangles, circles, diamonds, hexagons) carried out in technical triplicate. Statistical significance was calculated using one-way ANOVA with Dunnett’s multiple comparisons testing, \*\*\*\**p* ≤ 0.0001, ns not significant. (E) Quantification of whole cell GSL composition of i3Ns at 28 dpi by glycan profiling. The two shades of pink and blue reflect the two guides targeting each gene, GSL quantitation is displayed as percentage of total GSL abundance, mean ± SEM for N = 3-6 biological replicates. For clarity, statistical significance for individual pairwise GSLs comparisons are provided in Supplementary Figure S3. (F) Data from (E) grouped by lipid series into gangliosides (GM2, GD2, GM1a, GD1b, GD1a, GT1b, GQ1b), precursors (LacCer, GM3, GD3), o-series (GA2, GA1, GM1b, GD1α) and globosides (Gb3, Gb4, Gb5). Mean ± SEM. Significance calculated using two-way ANOVA with Dunnett’s multiple comparisons testing, \*\*\*\**p*≤0.0001, \*\*\**p*≤0.001, ns not significant.

Biosynthesis of GSLs involves multiple enzymes that function in a stepwise process to build glycan headgroups onto the ceramide backbone. Gangliosides are a subclass of GSLs containing sialic acid moieties and are particularly important in brain development and function. Two “gatekeeper” enzymes involved in the early steps of ganglioside synthesis are ST3GAL5 (GM3 synthase) and B4GALNT1 (GM2 synthase) with loss-of-function mutations causing severe childhood epilepsy[6] and an early-onset spastic paraplegia[7], respectively. Loss of either enzyme would remove all four predominant neuronal gangliosides (GM1a, GD1a, GD1b and GT1b, **Fig.1B**), but clinical severity differs in these diseases.

GM3 synthase deficiency (GM3SD) is an ultra-rare, autosomal recessive inherited metabolic disorder first characterized in the Old Order Amish community[6], where it was historically known as Amish Infantile Epilepsy Syndrome. However, it has been identified in several additional populations[8–10]. The clinical manifestation of GM3SD typically follows a severe and progressive course: while newborns often appear healthy, they rapidly develop refractory infantile-onset epilepsy, profound intellectual disability, and developmental stagnation within the first months of life[11]. Among the clinically validated pathogenic ST3GAL5 mutations[12], five have been evaluated for residual enzymatic activity. Activity assays conducted in HEK293T cells demonstrate that the most frequently reported variants (G342S, R288X, C195S, G201A and E355K) exhibit a complete lack of detectable enzymatic activity[13].

Hereditary spastic paraplegia 26 (HSP26) is another rare autosomal recessive neurological disorder of GSL synthesis caused by mutations in B4GALNT1. Clinically, HSP26 is distinguished from GM3SD by a later age of onset and a different progression: symptoms typically emerge in late childhood or adolescence as progressive lower-limb spasticity and weakness, sometimes accompanied by cognitive impairment and peripheral neuropathies[14–16]. Enzymatic activity assays demonstrate that the pathogenic B4GALNT1 variant D433A exhibits no detectable activity[15], whereas other variants of unknown significance display a range of activities from undetectable to near-wildtype levels[15,17].

Murine models have been developed to investigate GM3SD and HSP26, yet both illustrate the limitations of cross-species modeling of GSL disorders. ST3GAL5 knockout mice exhibit informative phenotypes including metabolic alterations[18], peripheral neuropathies[19] and impaired spatial memory[20]. However, they do not reproduce the defining clinical features of GM3SD, notably neurodevelopmental arrest and early-onset epilepsy. In contrast, B4GALNT1 knockout mice display clearer nervous system pathologies, including axonal degeneration and myelination defects[21] and develop progressive motor dysfunction with reflex, strength, coordination and gait abnormalities[22]. Nevertheless, across both disorders, mouse models do not consistently capture the full spectrum and timing of human disease, consistent with prior observations that species differences in GSL metabolism can influence neurological phenotypes[23]. Existing human-based models, such as patient-derived fibroblasts[24] and ST3GAL5 knockout HEK293 cells[25], are limited by their distinct cellular lineages resulting in lipid compositions that do not accurately reflect the neuronal tissue, making them less effective for functional investigations. Human neuronal models are therefore essential to understand the pathogenic mechanisms of GSL-related disorders.

Human iPSC-derived neurons offer an excellent isogenic inducible model to study the biochemical and electrophysiological changes when GSL metabolism is disturbed. Mature cortical glutamatergic neurons can be generated within 14 days following doxycycline-induced NGN2 expression, and they acquire functional maturity, including synchronized electrical activity, by day 28[26–28]. This i3N system has been successfully used to investigate molecular mechanisms driving several neurodevelopmental diseases including early onset gangliosidoses[28–30].

Here we define how distinct ganglioside repertoires support human neuronal function. We generated i3N models of GM3SD and HSP26 by knocking down the expression of genes *ST3GAL5* or *B4GALNT1*, respectively. Both models lose the predominant neuronal gangliosides, yet only GM3SD neurons fail to develop coordinated network activity, whereas HSP26 neurons maintain normal firing. Proteomic profiling of the plasma membrane reveals a mechanistic basis for this defect, with GM3SD neurons exhibiting significant loss of surface ion channels, GPCRs, and synaptic adhesion molecules, while the HSP26 membrane proteome remains largely intact. Together, these findings provide a mechanistic explanation for the distinct cellular and clinical phenotypes of GM3SD and HSP26, establishing a framework for dissecting how sialylated versus non sialylated GSLs shape neuronal maturation and function.

## RESULTS

### Loss of B4GALNT1 or ST3GAL5 cause severe changes to neuronal GSL profiles

Cellular disease models were generated using CRISPRi-mediated knock down (KD) of *B4GALNT1* or *ST3GAL5* gene expression. CRISPRi guides targeting these genes were introduced into the dCas9 derivative of the i3N stem cells[27]. For each gene, two KD cell lines using different guide sequences (**Table S1**) were made from the parental dCas9 line: ΔB4GALNT1-1 and −2 to model HSP26; and ΔST3GAL5-1 and −2 to model GM3SD. An additional cell line using a scrambled (SCRM) non-targeting guide in the same dCas9 background was used as a control. Quantitative PCR (qPCR) analysis of neurons confirmed efficient KD of *B4GALNT1* and *ST3GAL5* gene expression (**Fig.1C,D**). mRNA levels were below 2% of SCRM levels in all cell lines except for ΔST3GAL5-2, which retained 9.6% gene expression. Additional qPCR analysis of all cell lines demonstrated expression of neuronal markers (MAP2 and β3-tubulin) confirming that the disease model lines differentiate normally and express relevant markers of neuronal maturity (**Fig.S1**).

Historically, thin-layer chromatography (TLC) has been the primary technique for comparing GSL profiles[31]. However, TLC is limited by the absence of a universal buffer system compatible with all GSL species, meaning unexpected changes in the GSL profile can be missed. Additionally, it is intrinsically qualitative and so doesn’t allow accurate quantitation across samples. Quantification of individual GSL species from cells is difficult using mass spectrometry due to several glycan moieties possessing the same molecular mass and the heterogeneity of individual species due to ceramide backbone differences[32]. An alternative strategy to accurately quantify the abundance of different ganglioside headgroups involves enzymatically cleaving and labeling the glycan headgroups specifically from GSLs[33–35]. Briefly, lipids including GSLs are extracted from cell lysates, the ceramide tails of GSLs are cleaved using endoglycoceramidase and the glycan headgroups are fluorescently labelled with 2-anthranilic acid. The labelled headgroups are then separated using HPLC for quantification of individual peaks against known standards. To understand how the GSL profile was altered in the HSP26 and GM3SD disease models, this enzymatic approach was employed to quantify the abundance of all ganglioside species (**Fig.1E, S2,S3**). All four KD cell lines show significant loss of the major neuronal gangliosides including GM1a, GD1b, GD1a and GT1b, as well as the simpler gangliosides GM2 and GD2. Interestingly, the other changes to the GSL profile differed significantly between the two disease models.

Disruption of *B4GALNT1* expression results in an enormous accumulation of the precursor GSLs GM3 and GD3 (**Fig.1E,F**). As these are both direct substrates of B4GALNT1 in the metabolic pathway this is an expected consequence of this genetic disruption. Interestingly, another B4GALNT1 substrate, lactosylceramide (LacCer), also accumulates but to a lesser extent supporting that the presence of ST3GAL5 in this cell line efficiently converts LacCer to GM3. The ΔB4GALNT1 neurons therefore have a very simple GSL repertoire consisting primarily of only two sialylated GSLs, GM3 and GD3. Neither of these GSLs are normally abundant in healthy neurons as they are almost completely converted in the Golgi to more complex GSLs.

Loss of *ST3GAL5* expression also results in accumulation of high levels of the precursor lipid LacCer. Some of this precursor GSL is converted to the o-series GSLs GA2, GA1, GM1b and GD1α. These GSLs are not normally present in neurons, and the presence of B4GALNT1 in these cells has mediated production of this lipid class in the ΔST3GAL5 neurons. Surprisingly, there is also the appearance of globo-series GSLs: Gb3, Gb4 and Gb5 (also known as SSEA3). These GSLs are also not usually present in neurons, normally being enriched on non-neuronal cells including red blood cells and stem cells[36,37]. The enzyme A4GALT is responsible for the conversion of LacCer to Gb3[38]. *A4GALT* gene expression is very low in healthy neurons and in SCRM neurons explaining why no globo-series GSLs are normally present in neurons[27]. *A4GALT* expression was not upregulated in response to ST3GAL5 KD, supporting that the production of globo-GSLs is the result of abnormally increased levels of the LacCer substrate rather than gene dysregulation (**Fig.1G**). These analyses demonstrate that the GSL repertoire of the ΔST3GAL5 neurons has shifted away from the sialylated neuronal GSL species to a complex combination of different non-neuronal, primarily non-sialylated GSLs (**Fig.1F**).

### ST3GAL5 disruption severely impacts neuronal firing

To explore the impact of GSL repertoire changes on neuronal function, electrical activity was measured using a multi-electrode array (MEA). MEA allows for the non-destructive monitoring of spontaneous electrical signaling of neurons over time. Neurons were cultured on 24-well MEA plates and electrical signaling was monitored via an array of electrodes with each electrode measuring spikes of field potential. Electrical activity was measured from when neurons start becoming electrically active at around 14 days post-induction (dpi) until 35 dpi. The level of activity, measured as total number of spikes, increased over time in all cell lines but was severely and consistently reduced in the ΔST3GAL5 neurons (**Fig.2A**). This reduced spike count is clearly evident in the individual timecourse measurements taken at 28 dpi and 35 dpi (**Fig.2B,C** and **S4A,B**). The extent of this disruption in ΔST3GAL5 neurons varied across independently differentiated replicates (**Fig. S5,6**) but across the entire 35-day time period the number of spikes consistently failed to reach SCRM levels. This strong phenotype was not consistently observed in the ΔB4GALNT1 neurons (**Fig.2A** and **S4-6**) suggesting the ΔST3GAL5 neurons have a distinct defect in formation or propagation of electrical signals.

**Figure 2:**
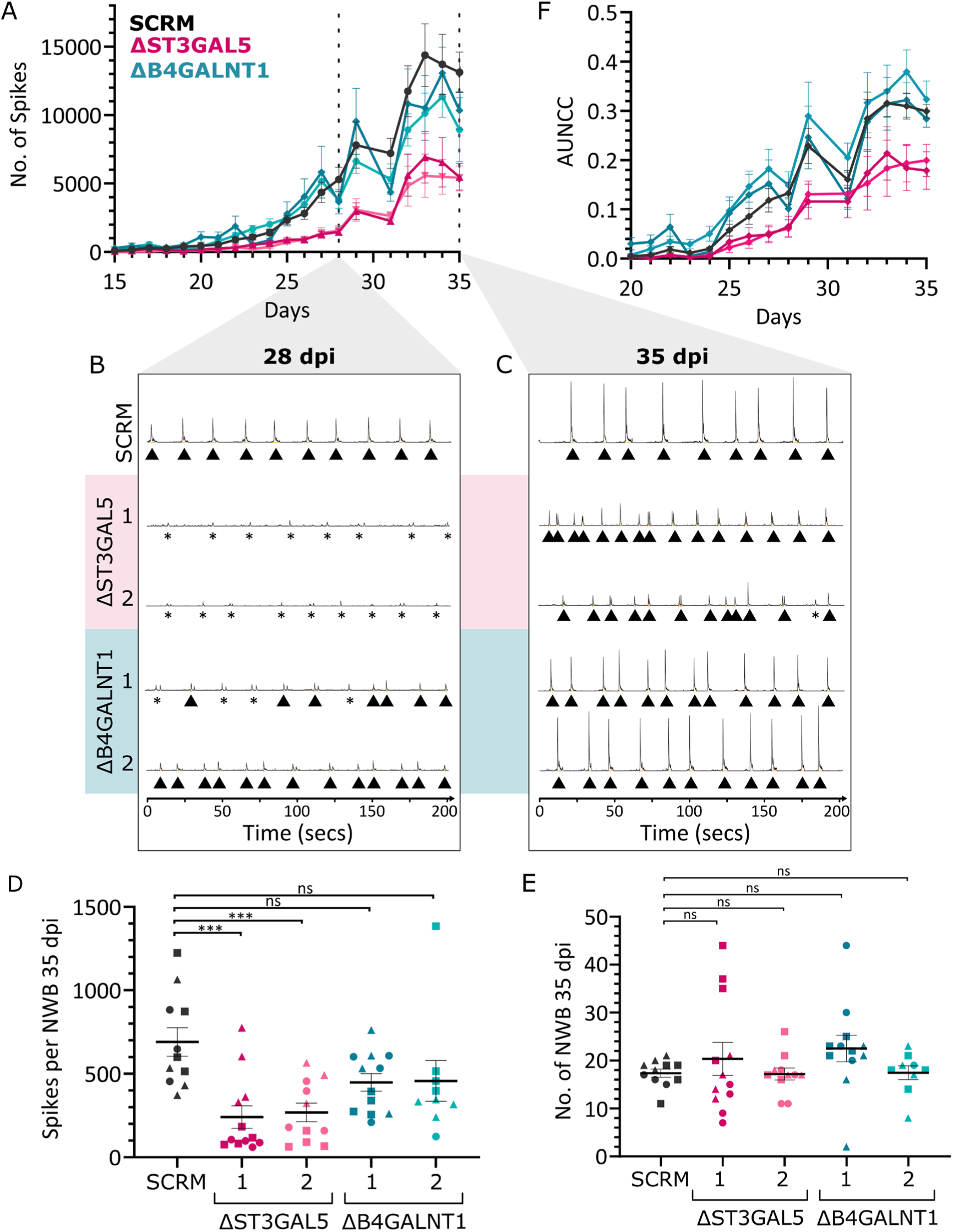
ΔST3GAL5 but not ΔB4GALNT1 neurons display severely disrupted electrical signaling. SCRM, ΔST3GAL5 and ΔB4GALNT1 neurons were cultured in 24 well multi-electrode array (MEA) plates and recordings were taken throughout differentiation. (A) Time courses monitoring number of spikes are shown for all cell lines with dashed lines indicating 28 and 35 dpi. The two shades of pink and blue reflect the two guides targeting each gene, Mean and SEM are shown for 9-12 wells measured per condition over N=3 biological replicates. Representative frequency histograms of spikes at 28 dpi (B) and 35 dpi (C). Triangles indicate network burst events. Asterisks represent synchronous firing events that don’t meet the threshold required to be a network burst. Spikes per NWB (D) and number of NWB (E) at 35 dpi for all cell lines. Mean ± SEM are shown for 9-12 wells measured per condition across N=3 biological replicates (circle, square, triangle). Significance calculated using one-way ANOVA with Dunnett’s multiple comparisons testing against SCRM, \*\*\**p*≤0.001, ns not significant. (F) Area under normalized cross-correlation (AUNCC) over time. Mean +/- SEM is shown for 9-12 wells measured per condition across N=3 biological replicates.

In addition to spike counts, MEA measurements can be analysed in terms of electrical bursts and network bursts (NWB). When >5 spikes were measured on a single electrode within 100 ms these were considered bursts, and when >35% of electrodes in a well registered bursts this was considered a NWB. SCRM neurons displayed strongly synchronized activity across the well from around 24 dpi with a regular pattern of NWB established from 28 dpi (**Fig.2B**, *triangles*, **Fig.S5-7**). Despite evidence for synchronous activity in ΔST3GAL5 neurons at 28 dpi, usually these did not meet the criteria to be a NWB due to the reduced number of spikes (**Fig.2B**, *asterisks*). As the electrical activity of ΔST3GAL5 neurons increased over time, these events crossed the threshold to be considered NWBs (**Fig.2C**) allowing for analysis of NWB behaviour at 35 dpi. Notably, the duration and number of spikes per NWB were significantly reduced in ΔST3GAL5 neurones (**Fig.2D, S4D-F**). However, no difference was seen in NWB frequency across cell lines (**Fig.2E**). ΔB4GALNT1 neurons display no significant reduction in mean duration or spikes per NWB (**Fig.2D, S4C-F**) although there was substantial variability, including several examples of very weak NWB behaviour (**Fig.2B**). The delayed and reduced electrical activity in ΔST3GAL5 neurons suggests the cells have impaired formation or function of synapses that is not seen in ΔB4GALNT1 neurons. In addition to probing the intensity of activity, MEA measurements also allow investigation into coordination of signaling through measurements of synchrony. The area under normalized cross correlation (AUNCC) measures how often and strongly neurons co-activate. The AUNCC is consistently reduced in the ΔST3GAL5 but not ΔB4GALNT1 neurons (**Fig.2F, S4G-H**). Taken together, this suggests the ΔST3GAL5 neurons have disrupted coordination of signaling in addition to reduced activity.

### Removal of globo- or o-series GSLs does not rescue the ST3GAL5 disrupted electrical firing

Gangliosides are the only highly abundant GSLs found in healthy neurons, and both ΔST3GAL5 and ΔB4GALNT1 neurons have large reductions in ganglioside abundance. However, only ΔST3GAL5 neurons have a severe electrical signaling phenotype. Therefore, either ΔB4GALNT1 neurons can better compensate for the loss of gangliosides or ΔST3GAL5 have an additional change in their GSL composition that is responsible for the disrupted electrical signaling. ΔST3GAL5 neurons have increased abundance of the non-neuronal globoside and o-series GSLs. To test whether disruption to electrical signaling in ΔST3GAL5 neurons is due to gain of globoside or o-series GSLs, double KD lines were generated.

To remove the globoside or o-series GSLs, guide RNAs were introduced into the ΔST3GAL5 background targeting the *A4GALT or B4GALNT1* genes respectively (**Fig. 3A**). qPCR of neurons at 28 dpi confirms efficient KD of *ST3GAL5* is maintained in all cell lines as well as efficient KD of additional genes in the respective double knockdowns (**Fig.3B**). GSL profiling confirms that in all KD lines significant reduction of ganglioside abundance is maintained or enhanced (**Fig. 3C**). Although KD of *B4GALNT1* expression in the ΔST3GAL5 background was the least efficient, at <15% of SCRM, it was still sufficient to reduce o-series (GA2, GA1, GM1b, GD1α) GSLs abundance back to near-normal levels (**Fig. 3C, S8**). Notably, in this double KD a single GSL, LacCer, now represents almost 80% of the total GSLs in these neurons. Although expression of A4GALT is already very low in neurons, KD of this gene in the ΔST3GAL5 background, to <5% of SCRM, reduces the globoside GSLs Gb3 and Gb4 back to normal levels (**Fig. 3C, S8**).

**Figure 3:**
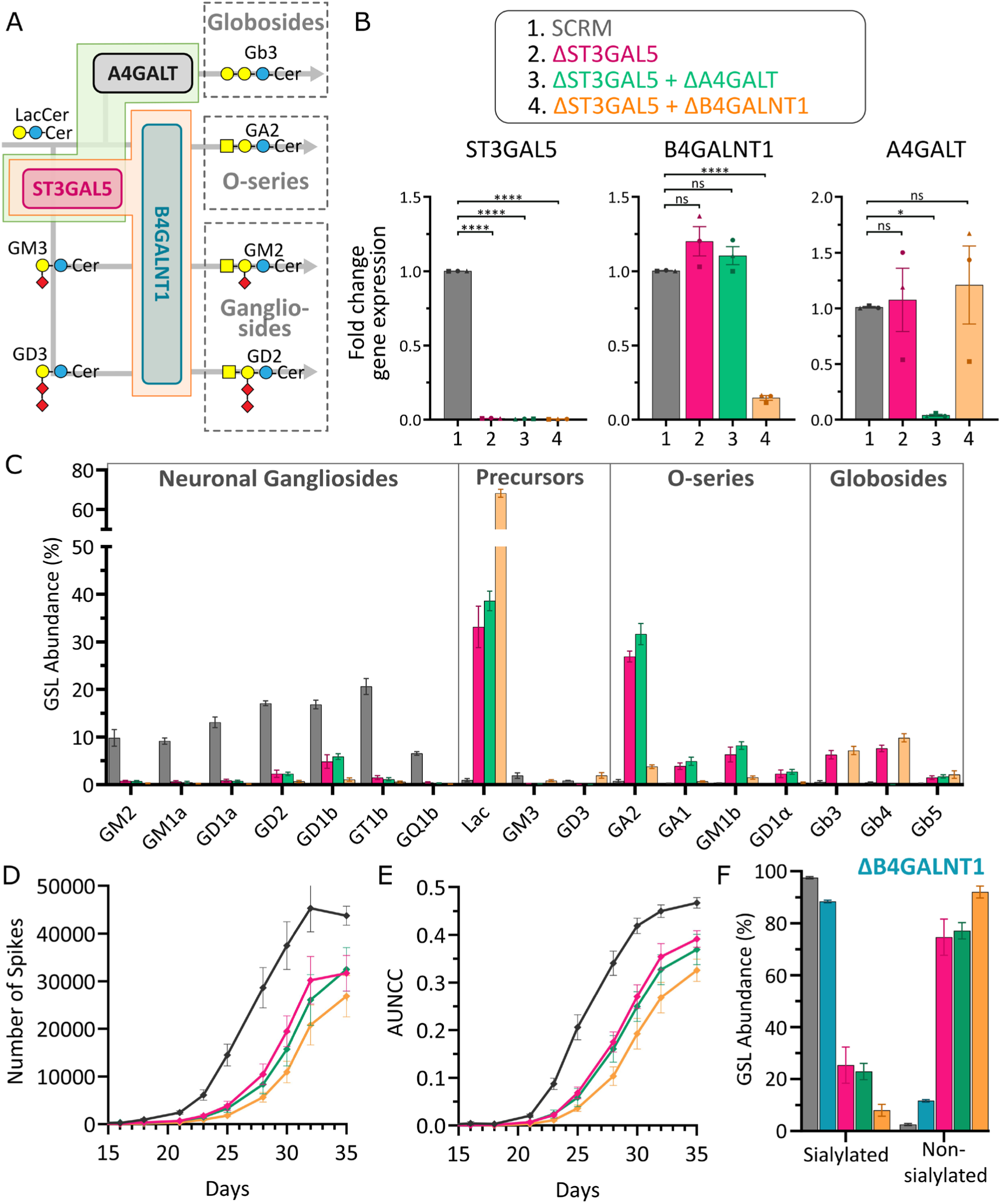
Accumulation of globosides or o-series lipids is not responsible for the disrupted electrical signaling in ΔST3GAL5 neurons. (A) Truncated schematic of GSL synthesis, highlighting the generation of double knock-down cell lines ΔST3GAL5+ΔA4GALT (green) and ΔST3GAL5+ΔB4GALNT1 (orange). Lipid series (dashed lines) and selected enzymes (solid lines) are boxed. (B) qPCR of i3N at 28 dpi, shown as fold change against SCRM. Mean ± SEM is shown for N=3 biological replicates (circle, square, triangle). Significance was calculated using one-way ANOVA with Dunnett’s multiple comparisons testing against SCRM, \*\*\*\**p*≤0.0001, * *p*≤0.05, ns not significant. (C) Quantification of whole cell GSL composition of i3Ns at 28 dpi by glycan profiling. GSL quantitation is displayed as percentage of total GSL abundance. Mean ± SEM for N=3 biological replicates. For clarity, statistical significance for individual pairwise GSLs comparisons are provided in Supplementary Figure S8. Time course of MEA data showing (D) number of spikes and (E) area under normalized cross-correlation (AUNCC) over time. Mean ± SEM are shown for 9-12 wells measured per condition across N=3 biological replicates. (F) Data from (Fig. 1E and 3C) grouped by sialic acid content into sialylated (GM2, GD2, GM1a, GD1b, GD1a, GT1b, GQ1b, GM3, GD3, GM1b, GD1α) or non-sialylated (LacCer, GA2, GA1, Gb3, Gb4, Gb5).

MEA analysis of electrical firing of these different compound KD lines demonstrates that loss of *A4GALT* or *B4GALNT1* expression and consequent removal of globoside or o-series GSLs, respectively, does not rescue the electrical signaling phenotype observed in ΔST3GAL5 neurons (**Fig.S9-11**). Number of spikes (**Fig.3D, S11D-E**), area under normalized cross correlation (**Fig.3E, S11F-G**), spikes per NWB and NWB duration (**Fig.S11A-B** and **H-K**) remain significantly reduced in all double KD lines compared to SCRM. This supports that the accumulation of globoside or o-series lipids are not the cause of the reduced electrical activity or disrupted coordination of signaling observed in ΔST3GAL5 neurons. Instead, it suggests that the precursors that accumulate in ΔB4GALNT1 neurons are better able to compensate for the loss of gangliosides than the globoside and o-series GSLs that accumulate in ΔST3GAL5 neurons. As LacCer accumulates in all KD lines including ΔB4GALNT1, it seems likely that GM3 and GD3 are the precursor GSLs responsible for this compensation (**Fig.1E** and **3C**). A notable common feature of GM3, GD3 and all the gangliosides that is not shared with the globoside or all o-series GSLs, is they contain a sialic acid moeity. Despite the relatively simple structure of these precursors, the GSL composition in ΔB4GALNT1 neurons retains a high abundance (>80%) of sialylated GSLs, similar to that of healthy neurons and distinct from that of the other KD models (**Fig. 3F**). Therefore, it is likely that this sialic acid moiety is important for why GD3 and GM3 can partially compensate for the loss of complex gangliosides, whereas the globoside and o-series lipids cannot. In further support of this hypothesis, the ΔST3GAL5+ΔB4GALNT1 neurons that possess the lowest abundance of sialylated GSLs also demonstrate the strongest electrical deficiencies (**Fig.3D-F, S11**).

### Loss of sialylated GSLs has a significant impact on the protein repertoire of the plasma membrane

Previous work has demonstrated that disruption of GSL metabolism can alter the protein composition of the PM[4,28,39]. Alterations to the PM proteome might thus explain the disruption to electrical firing in ΔST3GAL5 neurons. Importantly, comparison of the PM proteomes of the ΔST3GAL5 and ΔB4GALNT1 disease models allow for the dissection of which GSLs are most critical for protein delivery and stability at the PM, as well as which proteins could be contributing to the disrupted electrical phenotype. As SCRM neurons have formed mature functional synapses by 28 dpi but ΔST3GAL5 neurons have not established sustained NWBs (**Fig.2B**), this time point represents a particularly interesting stage in the development of network behaviour and was chosen for molecular analyses. Due to the relatively low abundance of individual membrane proteins, the PM composition was profiled using aminooxy-biotin labeling of intact cells and enrichment of proteins with streptavidin prior to mass spectrometry analysis. Proteomics datasets were filtered to remove non-PM proteins and data from the two individual guides for each KD were combined. High-confidence targets were selected based on a significant change (<0.05 adjusted *p*-value) in abundance of at least 1.65-fold compared to SCRM controls. Abundance changes are displayed as volcano plots and reveal substantial loss of several PM proteins in ΔST3GAL5 (**Fig.4A** and **Tables S2-3**) yet almost no changes in the ΔB4GALNT1 neurons (**Fig.4B** and **Tables S4-5**).

**Figure 4.**
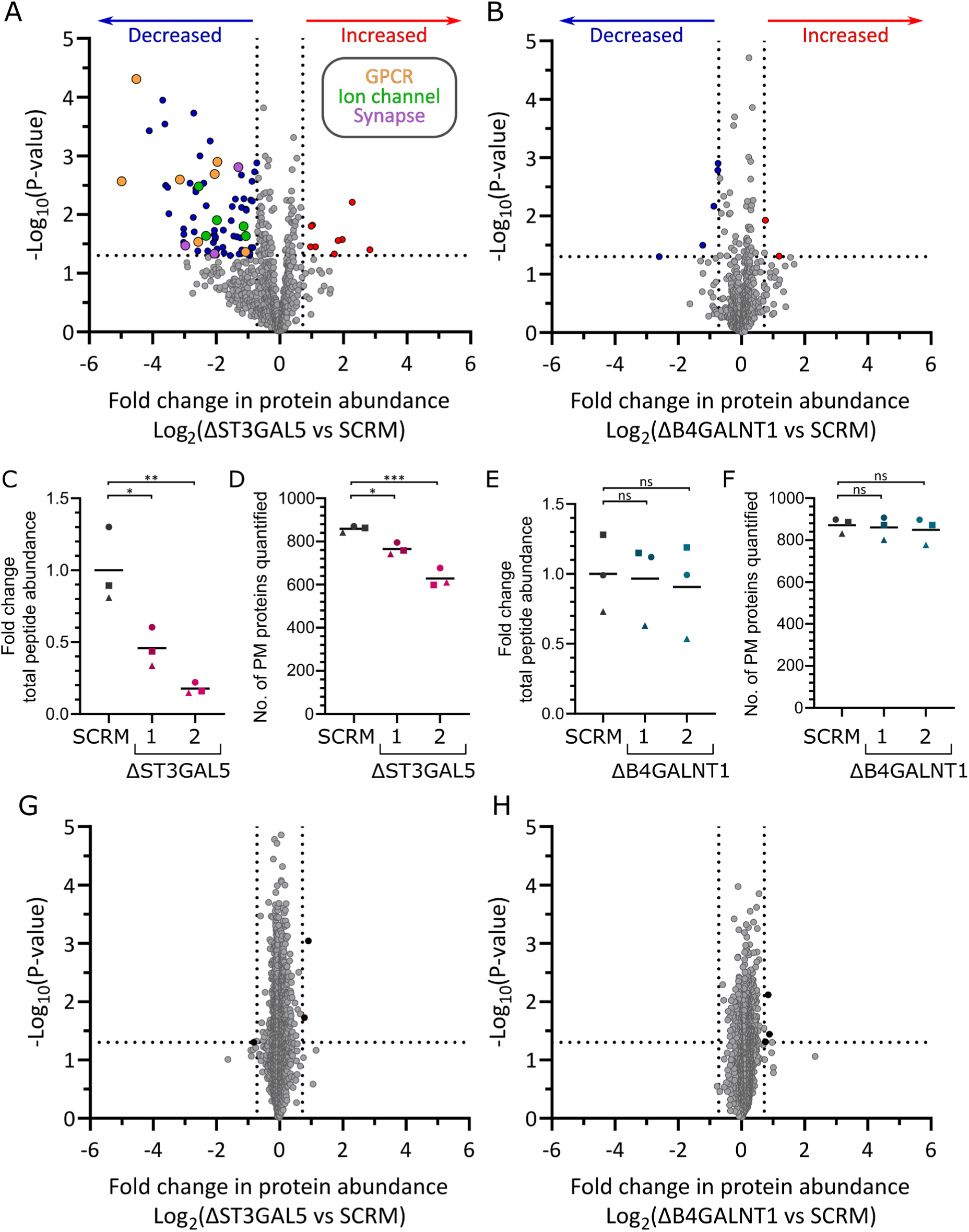
Loss of sialylated GSLs changes the neuronal cell surface proteome. Quantitative mass spectrometry following enrichment of PM proteins from ΔST3GAL5 and ΔB4GALNT1 neurons compared with the SCRM control. Plasma membrane proteomics data are displayed as volcano plots of ΔST3GAL5 (A) and ΔB4GALNT1 (B) neurons compared to SCRM neurons and shown with average fold change (x-axis) across three biological replicates and significance (y-axis, two-sided t test) across the three replicates. Protein targets are coloured based on reduced (blue) or increased (red) abundance with some specific protein classes labelled including GPCRs (orange), ion channels (green) and synaptic proteins (purple). (C) Total peptide abundance of ΔST3GAL5 samples, presented as fold change against SCRM. (D) Number of quantified plasma membrane proteins in ΔST3GAL5 samples. Equivalent analyses are shown for ΔB4GALNT1 samples in (E) and (F). Significance was calculated using one-way ANOVA with Dunnett’s multiple comparisons testing against SCRM, \*\*\**p*≤0.001, \*\**p*≤0.01 * *p*≤0.05, ns not significant. Volcano plots for whole cell proteomics data for ΔST3GAL5 (G) and ΔB4GALNT1 (H) neurons compared to SCRM neurons. Data displayed as for panels (A) and (B).

Interestingly, there was a decrease in the total abundance of peptide and number of quantified PM proteins in the ΔST3GAL5 samples compared to the SCRM despite normalization of the whole sample protein concentration prior to PM protein enrichment (**Fig.4C,D**). The drop in quantified proteins and peptide is not observed in ΔB4GALNT1 neurons (**Fig.4E,F**) suggesting that the loss of sialylated GSLs has significantly altered the capacity of ΔST3GAL5 neurons to deliver and/or retain membrane proteins at the PM. A significant drop in peptide or protein abundance is not seen in the whole-cell proteome of ΔST3GAL5 or ΔB4GALNT1 neurons and very few proteins are significantly changed in abundance in either cell line compared to SCRM (**Fig.4G,H** and **Tables S6-8**). This suggests that observed changes in protein abundance at the PM are caused by mislocalization of proteins rather than overall changes in protein expression, synthesis or stability.

Due to the large-scale remodeling of the plasma membrane proteome it is highly likely that no single change is the key determinant of disrupted electrical signaling or ST3GAL5-associated disease, and that this is instead a complex consequence of the large number of proteins lost from the PM. Proteins that may be directly contributing to disrupted electrical activity include voltage- and ligand-gated ion channels, which are essential for synaptic function and electrical signaling. Five ion channels are significantly downregulated from the PM in ΔST3GAL5 neurons (HCN2, KCNH7, KCNA3, CLCN4 and HTR3A). The voltage-gated, non-selective cation channel HCN2 is important for controling conduction velocity, setting the resting membrane potential and ensuring rhythmic firing of neurons[40–42]. Voltage-gated Na^+^ channels enable rapid depolarization during action potential and voltage-gated K^+^ channels such as KCNH7 (a.k.a. ERG3) and KCNA3 (a.k.a. Kv1.3) are required to stabilize the resting potential and repolarize the membrane[43,44]. Ligand-gated channels such as the serotonin-activated HTR3A mediate rapid synaptic transmission in response to neurotransmitters[45]. CLCN4 (a.k.a. CIC-4) is a predominantly intracellular Cl^-^ channel but has been identified at the PM of neurons[46]. All five of these channels have been associated with epilepsy and other neurological diseases[44,47–51].

Another group of proteins that could contribute to ST3GAL5-associated disease are G-protein coupled receptors (GPCRs), which are a diverse class of membrane receptors that are localised to GSL-rich membrane microdomains and are activated by extracellular stimuli such as neurotransmitters. Seven GPCRs (CHRM4, CXCR4, KISS1R, NPY1R, OPRL1, PTGER3, TRHR) are significantly downregulated from the PM in ΔST3GAL5 neurons. Activation of GPCRs can influence effector proteins and the production of second messengers such as cAMP which is crucial for the function of pre- and post-synaptic plasticity. GPCRs play key roles in regulating ion channel activity and neurotransmitter release and so a loss in their abundance at the cell surface is likely to disrupt electrical signaling. Although the activity of some of these GPCRs may not be triggered in a monoculture of cortical glutamatergic neurons, the reduced abundance of these at the PM remains relevant for ST3GAL5-associated disease.

Transmission of electrical and chemical signals across synapses, mediated in part by ion channels and GPCRs requires coordinated formation and organization of synapses. Pre- and post-synaptic adhesion molecules such as NRXN3 and LRRTM3, both of which are downregulated in ΔST3GAL5 neurons play major roles in modulating this organization as well as influencing synaptic plasticity[52–55].

A small number of proteins appear upregulated at the surface of ΔST3GAL5 neurons (**Fig. 4A**). However, most of these were quantified in SCRM neurons but not in ΔST3GAL5 neurons and the appearance of them being upregulated is likely an artefact of imputation (**Fig. S12**).

## DISCUSSION

Comparative analysis of human iPSC-derived neuronal models of ST3GAL5 and B4GALNT1 deficiency reveals a direct relationship between GSL composition and cellular function. By expanding these models into dual knockdowns, we clearly demonstrate that it is the loss of sialylated neuronal gangliosides, rather than the gain of non-neuronal GSLs, that disrupts propagation and coordination of electrical signaling. We further demonstrate that the synthesis of simple sialylated GSLs can prevent this signaling defect, highlighting the essential role of lipid sialylation for neuronal function. Mechanistic insights into disease pathology are demonstrated by the contrasting PM proteome profiles of these two models of distinct GSL metabolic defects. Absence of sialylated GSLs in the ΔST3GAL5 neurons causes widespread depletion of PM proteins, several of which are crucial for neuronal functions. This loss explains the electrophysiological phenotype in ΔST3GAL5 neurons and allows for a mechanistic understanding of the severe neurological phenotypes in GM3SD patients. Importantly, the presence of simple sialylated GSLs in ΔB4GALNT1 neurons can, to some extent, compensate for this loss as demonstrated by almost wildtype electrical firing and an essentially unchanged PM proteome.

While ΔST3GAL5 mouse models of GM3SD display metabolic and behavioural changes, they lack the microcephaly, severe neurodevelopmental delay and epilepsy characteristic of the human condition[18,56,57]. This is unlikely to reflect a reduced dependence on gangliosides, as ΔST3GAL5+ΔB4GALNT1 mice develop severe post-weaning neurodegeneration and die at 2-3 months, supporting that gangliosides are essential for the higher order development of the mouse CNS[58,59]. Instead, it suggests loss of function mutations in ST3GAL5 differentially affect the GSL profile of mice and humans, resulting in different severities of disease. Across ΔST3GAL5 mouse brain[18,57,60], patient samples[6,8,9,11,61], and our iPSC-derived neurons, gangliosides and sialylated precursors (GD3 and GM3) are consistently reduced. In contrast, patient samples[6,9,61] and our iPSC models accumulate globosides and non-sialylated o-series lipids (GA2, GA1), changes not observed in mice. However, compound knockdowns demonstrate that these non-neuronal GSLs did not drive the electrophysiological phenotype. The important difference is that the human ΔST3GAL5 GSL profile is dominated by non-sialylated lipids, whereas the dominant species in the brains of ΔST3GAL5 mice are the sialylated o-series lipids GM1b and GD1α[6,8,9,11,18,57,60,61]. GD1α may be particularly important, as it is largely absent in human samples but retained in mice and remains partially sialylated. This, in combination with a greater proportion of GM1b could be sufficient to partially compensate and reduce the severity of disease in mice. Alternatively, mice may express additional compensatory GSL species that remain undetected due to limitations of thin-layer chromatography, which could be resolved using quantitative glycan profiling approaches.

Unlike ST3GAL5, loss of function mutations in B4GALNT1 have comparable consequences in mice and humans, including gait defects and peripheral axonal degeneration[7,16,21,22,62]. This likely reflects shared alterations in the GSL profile, with decreased gangliosides and increased GM3 and GD3 observed in both mice[62–66] and in our ΔB4GALNT1 neurons. The PM proteome (**Fig.4**) and electrical firing (**Fig.2&3**) of ΔB4GALNT1 neurons remain largely intact, despite neurodegenerative disease in patients and mice, suggesting that GM3 and GD3 provide only partial or transient compensation for the loss of gangliosides. Consistent with this, HSP26 shows delayed childhood or adolescent onset, whereas GM3SD presents in early infancy with severe epilepsy[7,9,67]. GM3 and GD3 may support initial protein delivery to the cell surface but fail to maintain long-term surface localization or stable membrane microdomains. GD3 appears particularly important, as the majority of mice expressing only GM3 die before 30 weeks of age[68,69].

The question remains as to why the loss of the sialylated GSLs has such a profound impact on the PM proteome. Are these proteins not delivered to the PM or are they being prematurely internalized? Extensive evidence supports the role of GSLs in biosynthetic sorting in the late secretory pathway, including recycling of membrane proteins to and from the PM, but specific details for how different GSLs contribute to sorting remains elusive[70–75]. There is also growing evidence supporting the importance of sialylated glycans on proteins and lipids for modulating endocytosis, with glycan-binding galectins playing crucial roles in tuning endocytic processes[76]. Glycolipid-lectin-driven endocytosis is a clathrin-independent mechanism for membrane protein internalization, with the clustering of GSL-bound oligomeric lectins inducing membrane bending to form endocytic pits[76–79]. However, the relevant glycan determinants, cargo specificity, and molecular mechanisms remain unresolved[76,80]. Studies of viral and toxin entry further suggest that sialic acid positioning on GSLs influences intracellular trafficking routes[81], although these mechanisms are difficult to dissect due to the promiscuous GSL binding of pathogen-related proteins[82–85].

Our observations, here and previously, show that distinct subsets of proteins change in PM abundance depending on different alterations in the cellular GSL repertoire[28,39]. This supports specificity in the disruption rather than a general impact on membrane microdomain structures or membrane dynamics. Such specificity may arise from direct interactions between protein extracellular domains (ECDs) and GSL headgroups. Several proteins have been identified as GSL-binders[86–88], and binding to distinct headgroups can functionally modulate the activity of some membrane proteins[89–91]. However, their specificity is difficult to resolve as complex GSLs intrinsically contain simpler glycan motifs that may enable protein binding, albeit more weakly than to the complex headgroup. This is consistent with our observation that GM3 and GD3 can partially restore membrane trafficking while leaving cells susceptible to progressive degeneration.

Our data support that GM3SD is not only a lipid metabolic disorder but also a secondary channelopathy/synaptopathy driven by lipid-dependent protein mislocalization. Interpretation of functional phenotypes is complicated by the absence of benchmarking MEA datasets distinguishing developmental delay from primary epileptic network instability, making it difficult to assign the failure of ΔST3GAL5 neurons to establish sustained network bursts to either mechanism alone. Although the observed phenotypes in ΔST3GAL5 neurons likely arise from the combined loss of multiple PM proteins, examining the known roles of these proteins in neurological disorders offers functional insights into the pathogenic drivers. Three protein classes emerge as likely contributors: ion channels, GPCRs, and adhesion molecules. Among the downregulated ion channels, HTR3A[92] and HCN2[93] affect neuronal depolarization, with HCN2 implicated in both neurodevelopmental disorders[48] and epileptogenesis[49,94]. HCN channels exert complex effects on neuronal excitability, as they exhibit a “dual nature”: while their opening depolarizes the membrane and increases excitability, it also decreases membrane resistance, which dampens excitability[40]. Conversely, the voltage-gated potassium channels KCNA3 and KCNH7 are essential for repolarization[43,44], and linked to epileptic and developmental disorders[44,50,51]. Together, these changes indicate that both depolarization and repolarization processes may be profoundly disrupted in ΔST3GAL5 neurons.

In addition to ion channels that directly affect neuronal firing, changes in GPCRs and adhesion molecules represent a more nuanced layer of disruption. One particularly relevant example is the downregulation of KISS1R and NPY1R, the two most significantly depleted GPCRs in our dataset. Both receptors regulate intracellular cAMP levels[95,96], a second messenger that modulates HCN channel activity[97]. The therapeutic effects of their ligands, kisspeptin[98] and neuropeptide Y[99], in epilepsy further underscore the relevance of these GPCRs in general seizures. However, it is worth noting that several drugs that are used for treating epilepsies do not provide long-term relief in GM3SD, suggesting that the therapeutic targets of these small molecules may themselves be downregulated from the PM[100].

Intriguingly, several downregulated proteins in our PMP dataset, including LRRTM3[101], CXCR4[102] and HCN2[103], are associated with late-onset neurodegenerative pathologies such as Alzheimer’s disease. Furthermore, neuronal models of early-onset neurodegenerative lysosomal storage disorders (LSDs) driven by ganglioside accumulation, exhibit PM protein remodeling including depletion of CXCR4 and NRXN3[28]. Given the involvement of membrane microdomains in Alzheimer’s[104], Parkinson’s disease[105] and LSDs[106–108], PM protein depletion driven by GSL alterations may be a shared mechanistic link disrupting microdomain-dependent proteostasis.

The extensive remodeling of the plasma membrane proteome in ΔST3GAL5 neurons reveals a highly interconnected pathological landscape, in which the loss of sialylated gangliosides destabilizes multiple functionally interdependent protein networks. Rather than a single dominant defect, the resulting phenotype reflects a systems-level failure of membrane organization. This complexity suggests that restoring individual downstream pathways is unlikely to be sufficient. Instead, our findings point to sialylated gangliosides as central organizational elements required to maintain plasma membrane integrity and neuronal function.

## MATERIALS AND METHODS

### Cell line maintenance

Human CRISPRi-i3N stem cells[26] carrying a stably integrated dCas9 construct were cultured at 37 °C and 5% CO_2_ in complete E8 medium (Gibco™), with daily medium changes. Cells were grown on Matrigel-coated plates (1:50 dilution in DMEM/F-12 HEPES; Gibco™). For passaging, cells were detached either with 0.5 mM EDTA and re-plated as colonies in E8 medium, or with Accutase and re-plated as single cells in E8 medium supplemented with 50 nM chroman-1 (Tocris).

Differentiation of stem cells into neurons was performed as previously described, with minor modifications[26]. Following Accutase dissociation, 7 × 10^6^ or 1.5 × 10^7^ stem cells were seeded onto 10 or 15-cm Matrigel-coated dishes respectively. Cells were seeded in induction medium (IM), consisting of DMEM/F-12 HEPES supplemented with 1× N2, 1× NEAA, 1× GlutaMAX, and 2 μg/ml doxycycline. During initial plating, IM was additionally supplemented with 50 nM chroman-1, and the medium was replaced daily for the first three days.

Partially differentiated neurons were dissociated with Accutase and re-plated onto dishes coated with 100 μg/ml poly-L-ornithine (PLO) in cortical neuron (CN) medium, consisting of Neurobasal Plus supplemented with 1× B27, 10 ng/ml NT-3 (PeproTech), 10 ng/ml BDNF (PeproTech), and 1 μg/ml laminin. During re-plating, CN medium was supplemented with 1 μg/ml doxycycline. From this point onward, half-medium changes were performed three times per week.

### Generation of CRISPR knockdown cell lines

Guide RNAs targeting ST3GAL5, B4GALNT1 and A4GALT were designed using Dharmacon or the CRISPick server (Broad Institute; **Table S1**), selecting sites at or immediately upstream of the first exon. For targeting of a single gene, complementary oligonucleotides with appropriate overhangs were annealed and cloned into the BpiI site of the pKLV backbone (gift from the Prof. Evan Reid, Cambridge). For targeting a second gene, oligonucleotides were cloned into the BsmB1 site of the lentiCRISPRv2 backbone with a hygromycin resistance cassette (gift from Prof. Evan Reid, Cambridge)[109].

For lentiviral production, HEK293T cells were cultured in DMEM supplemented with 10% FBS at 37 °C and 5% CO_2_ and passaged twice weekly using Trypsin-EDTA. A total of 2 × 10^6^ cells were transfected in OptiMEM with 8 µl TransIT and 2 µg of plasmid DNA at a 3:2:1 ratio of CRISPRi pKLV construct:pCMVΔ8.91:pMD.G. The transfection mixture was added dropwise, and after 24 h the medium was replaced with E8. At 48 h post-transfection, the virus-containing E8 medium was collected, 0.45μm filtered to remove debris and stored at −70 °C.

5 × 10^5^ i3N stem cells were transduced with 2 ml of a 1:1 mixture of complete E8 medium and lentiviral supernatant supplemented with 50 nM chroman-1 and 10 µg/ml polybrene. Cells were seeded on Matrigel-coated 6-well plates and then transferred to 37 °C for incubation. After 24 h, the medium was replaced with fresh E8. At 48 h post-transduction, cells were switched to E8 medium containing 1 µg/ml puromycin and if targeting a second gene also 50 µg/ml hygromycin. Media was refreshed daily for at least 7 days. Surviving cells were subsequently maintained in puromycin and hygromycin-free E8 medium.

### Quantitative PCR (qPCR)

RNA was extracted from 2 × 10^6^ cells using the PureLink RNA extraction kit (Invitrogen) following the manufacturer’s protocol. cDNA was generated from purified RNA using the High-Capacity RNA-to-cDNA kit (Applied Biosystems) following the manufacturer’s protocol. qPCR reactions contained 1× TaqMan Fast Advanced Master Mix (Applied Biosystems), 50 ng cDNA, and 1× FAM-labelled TaqMan Gene Expression Assays (Applied Biosystems) specific to each target gene (**Table S9**). Reactions were run on a CFX96 Real-Time System (Bio-Rad), starting with 10 mins at 95 °C, followed by 40 cycles of 95 °C for 15 s and 60 °C for 1 min. Each experiment included N=3 technical replicates per gene and N=3 independent biological replicates. Gene expression changes were quantified using the ΔΔCt method with GAPDH as the reference gene[110].

### Quantification of gangliosides by HPLC

Glycosphingolipids were analysed as previously described[34,35]. Briefly, lipids were extracted from 2 × 10^6^ cells using a 4:8:3 chloroform:methanol:water mixture and purified via C18 solid-phase extraction (Telos, Kinesis). Bound lipid species were eluted twice with 1 mL 98:2 chloroform:methanol, followed by two 1 mL elutions of 1:3 chloroform:methanol and a final 1 mL methanol wash. Eluates were dried under a stream of nitrogen gas and digested overnight with recombinant endoglycoceramidase I (gift from the Platt Lab, Oxford). Released glycans were labelled with 2-anthranilic acid (2AA) and purified using DPA-6S amide SPE columns (Supelco). Purified 2AA-labelled glycans were separated and quantified by normal-phase HPLC. Peak areas were compared with a 2.5 pmol 2AA-labelled chitotriose standard (Ludger) to calculate absolute values, and values were presented as percentage GSL abundance for ease of comparison between samples. Glycan species were identified by matching retention times to a reference standard of commercially available gangliosides. Chromatographic separation was performed on a TSKgel Amide-80 column (Anachem) using a Waters Alliance 2695 separations module with an in-line Waters 2475 multi λ-fluorescence detector (excitation 360 nm, emission 425 nm). All runs were conducted at 30 °C using solvent compositions and gradient conditions described previously[34,35].

### Proteomics

#### Proteomics cell culture

Cell lines were differentiated in triplicate to generate three independent biological replicates. Partially differentiated 3 dpi i3N cultures of SCRM, ΔST3GAL5-1 and ΔST3GAL5-2 or SCRM, ΔB4GALNT1-1, and ΔB4GALNT1-2 were seeded onto PLO-coated plates. For whole-cell proteomics (WCP) cells were seeded at 2 × 10^6^ cells per well of a 6-well plate and for plasma membrane proteomics (PMP) at 1 × 10^7^ cells per 10-cm dish. Cultures were maintained until 28 dpi with half-medium changes performed thrice weekly.

#### Sample preparation for WCP

Cells were washed three times with ice-cold PBS and scraped into low-bind Eppendorf tubes. Pellets were collected at 500 × g for 10 min, PBS was removed, and samples were snap-frozen in liquid nitrogen and stored at −70 °C until simultaneous processing of biological replicates. Cell pellets were resuspended and lysed in 6 M guanidine hydrochloride/50 mM HEPES, pH 8.5. Samples were vortexed extensively, sonicated in a cold water-bath sonicator using five 30 s pulses, and clarified by centrifugation at 13,000 × g for 10 min at 4 °C. The supernatant was transferred to a fresh tube and centrifuged again under the same conditions. Proteins were reduced with DTT (final concentration 5 mM) for 20-30 min shaking at 56 °C, alkylated with iodoacetamide (final concentration 14 mM) for 20-30 min shaking at room temperature in the dark, and excess iodoacetamide was quenched by addition of further DTT (final concentration 9 mM) followed by incubation for 15 min at room temperature. Samples were then diluted with 200 mM HEPES, pH 8.5 to reduce the guanidine concentration to 1.5 M and digested with LysC for 3 h at room temperature. Samples were subsequently diluted further to 0.5 M guanidine with 200 mM HEPES, pH 8.5, supplemented with trypsin to a final concentration of 3.33 ng/μL, and incubated shaking overnight at 37 °C. The following day, digests were acidified with formic acid (final concentration 2.5% v/v) and centrifuged at 21,000 × g for 10 min, and the clarified supernatant was transferred to a fresh tube for peptide clean-up. Peptides were purified using 50 mg C18 Sep-Pak cartridges that were activated with acetonitrile and then formic acid-containing buffer (70% acetonitrile, 1% formic acid) before equilibration into 1% formic acid. After sample loading, cartridges were washed with 1% formic acid and peptides were eluted with formic acid-containing buffer (70% acetonitrile, 1% formic acid)

Eluates were dried by vacuum centrifugation and resuspended in 200 mM HEPES or 150 mM TEAB, pH 8.5. Peptide concentration was determined by micro-BCA assay, and 25-40 μg of peptide were taken forward for TMT labeling. For labeling, samples were adjusted to 40 μL in 200 mM HEPES, pH 8.5 and to a final acetonitrile concentration of 30%, then reacted with 5 μL TMT reagent (18.6 μg/ μL in anhydrous acetonitrile) for 1 h at room temperature. Reactions were quenched by addition of 5% hydroxylamine to a final concentration of approximately 0.5% and incubated for 15 min, after which samples were acidified to approximately 1% formic acid. Labelled samples were then combined in equal peptide amounts, and residual free tag was removed by a further Sep-Pak clean-up. Briefly, the Sep-Pak cartridge was wetted with 1 mL of 100% methanol, then by 1 mL acetonitrile, equilibrated with 1 mL of 0.1% trifluoracetic acid and the sample was loaded slowly. The cartridge was washed 3x with 1 mL trifluoracetic acid before eluting sequentially with 250 μL of 40% acetonitrile, 70% acetonitrile and 80% acetonitrile, before being dried in a vacuum centrifuge.

The peptides were fractionated by high-pH reverse-phase chromatography using an Ultimate 3000 ultra-high-performance chromatography (UHPLC) system (Thermo Scientific) equipped with a Kinetex Evo C18 (1.7 μm, 2.1 × 150 mm) column (Phenomenex). The mobile phases used were, HPLC-grade water 3:97(v/v) (eluent A); 100% LCMS-grade acetonitrile (eluent B) and 200 mM ammonium formate in HPLC-grade water, pH 10 (eluent C). Eluent C was maintained at 10%, while A and B were altered over the fractionation gradient. The elution gradient was 0-19% B in 10 min; 19-34% B in 14.25 min and 34-50% B in 8.75 min, followed by a 10 min wash of 90% B. Flow was set at 0.20 mL/min for the first 5 min while peptides which were reconstituted in 40 μL of eluent C were loaded onto the column. Thereafter the flowrate was increased to 0.40 mL/min for the rest of the protocol. 168 fractions were collected every 0.25 min between 10 and 52 min of the fractionation run. The fractions were concatenated into 12 pools and reduced to dryness in a refrigerated vacuum centrifuge.

#### Sample preparation for PMP

Sialic acids on the extracellular surface of plasma-membrane proteins were oxidized with sodium periodate to generate aldehydes, followed by aniline-catalysed oxime ligation with aminooxy-biotin. For biotinylation, 10 cm dishes of cells were washed three times with 5 ml ice-cold PBS (pH 7.4) and incubated in 5 ml biotinylation solution (1 mM sodium periodate, 100 μM aminooxy-biotin, 10 mM aniline in ice-cold PBS pH 6.7). Plates were wrapped in foil and rocked at 4 °C for 30 min. Reactions were quenched with 5 ml 2 mM glycerol, glycerol was removed, and cells were washed three times and then scraped into ice-cold PBS pH 7.4. Pellets were collected at 500 × g for 10 min, PBS was removed, and samples were snap-frozen in liquid nitrogen and stored at −70 °C until simultaneous processing of biological replicates. Pellets were lysed in 500 μl lysis buffer (10 mM Tris pH 7.4, 1% Triton X-100, 150 mM NaCl, 5 mM EDTA, protease inhibitors) in low-bind tubes. Lysis was performed by end-over-end rotation for 1.5 h at 4 °C. Lysates were cleared by centrifugation at 20,000 × g for 10 min, and supernatants were transferred to fresh low-bind tubes before quantification by BCA assay. Protein concentrations were equalized using lysis buffer so that all samples contributed identical total protein mass to enrichment. High Capacity NeutrAvidin Agarose Resin (Thermo Scientific, 50 μL slurry per sample) was washed four times with 1 ml lysis buffer (resuspension, 500 × g for 5 min, aspiration, repeat). Normalized lysates were added to the washed beads and incubated by end-over-end rotation for 2.5 h at 4 °C. For washing, bead-lysate mixtures were transferred onto SnapCap filter columns (Pierce) mounted on a vacuum manifold. After draining, beads were washed sequentially with 20 × 400 μL lysis buffer, 20 × 400 μL 0.5% SDS in PBS pH 7.4, and 10 × 400 μL urea buffer (6 M urea, 50 mM TEAB, pH 8.5). Columns were removed, capped, and beads were incubated with 400 μL reduction/alkylation solution (10 mM TCEP, 20 mM iodoacetamide in urea buffer) for 30 min at room temperature in the dark with shaking at 850 rpm. Columns were returned to the manifold, drained, and washed an additional 10x with urea buffer. Beads were then resuspended in 400 μL 50 mM TEAB and transferred to low-adhesion tubes. Columns were rinsed twice more with 400 μL TEAB and the rinses were pooled with the bead suspension. Beads were pelleted at 500 × g for 2 min and supernatant was removed. Beads were resuspended in 50 μL of 50 mM TEAB containing 0.5 μg MS-grade trypsin (Pierce; stock 1 μg/μL in 50 mM acetic acid) and digested overnight at 37 °C with shaking at 850 rpm. The next day, beads were pelleted at 500 × g for 5 min, and the supernatant was collected. Beads were washed with an additional 40 μL of 50 mM TEAB; the wash was pooled with the digest supernatant. Eluates were dried using a vacuum centrifuge and stored at −70 °C.

#### Mass Spectrometry for WCP samples

WCP samples were analysed using tandem mass tag (TMT) labeling and data-dependent acquisition (DDA). Mass spectrometry data for WCP were acquired by Dr. Robin Antrobus of the CIMR proteomics facility on an Orbitrap Lumos instrument as previously described[111]. Peptides were separated on an RSLC3000 nano-UPLC system equipped with a 300 μm × 5 mm Acclaim PepMap μ-precolumn and a 75 μm × 50 cm, 2.1-μm particle Acclaim PepMap RSLC analytical column. The loading solvent was 0.1% formic acid (FA), and the analytical solvents were 0.1% FA (solvent A) and 80% acetonitrile with 0.1% FA (solvent B). All separations were performed at 40 °C, with samples loaded at 5 μl/min for 5 min before application of one of two gradients: 3-7% B over 4 min and 7-37% B over 173 min followed by a 4-min wash at 95% B and 15-min re-equilibration at 3% B, or 3-7% B over 3 min and 7-37% B over 116 min followed by identical wash and re-equilibration. Quantification used a MultiNotch MS3-based TMT workflow[112], with MS1 acquired from m/z 380-1,500 at 120,000 resolution (2 × 10^5^ AGC target, 50-ms maximum injection time), MS2 acquired in the ion trap following quadrupole isolation (m/z 0.7), CID fragmentation (NCE 34), turbo-scan mode (1.5 × 10^4^ AGC target, 120-ms maximum injection time), and MS3 performed in synchronous precursor selection mode selecting the top 10 MS2 ions for HCD fragmentation (NCE 45) and Orbitrap detection at 60,000 resolution (1 × 10^5^ AGC target, 150-ms maximum injection time). Parallelisable time was disabled, and the full MS1/MS2/MS3 cycle time was 3 s. Mass spectrometry data have been deposited with the ProteomeXchange Consortium via the PRIDE repository with the dataset identifier PXD076397.

#### Mass spectrometry for PMP samples

PMP samples were analysed using label-free data-independent acquisition (DIA) mass spectrometry. Mass spectrometry data for PMP were acquired by Dr. Robin Antrobus of the CIMR proteomics facility on a Thermo Q Exactive Plus mass spectrometer (Thermo Scientific) equipped with an EASYspray source and coupled to an RSLC3000 nano-UPLC system. Digested peptides were separated on a 50 cm C18 PepMap EASYspray column maintained at 40 °C at a flow rate of 300 nL/min. A gradient was run using solvent A (0.1% formic acid) and solvent B (80% acetonitrile, 0.1% formic acid), increasing from 3% to 40% solvent B over 90 min, followed by a 5 min wash at 95% solvent B. Full MS spectra were acquired at a resolution of 35,000 across an m/z range of 350-1430, and MS/MS spectra were collected in DIA mode. Fragmentation spectra were acquired at 17,500 FWHM following HCD, using a 30 m/z isolation window, an AGC target of 1 × 10^6^, and a loop count of 18. Mass spectrometry data have been deposited with the ProteomeXchange Consortium via the PRIDE repository with the dataset identifier PXD076397.

#### Analysis of mass spectrometry data

WCP data was processed in Proteome Discoverer 2.2 using the Sequest search engine and PMP data was processed with default parameters in DIA-NN version 2.2.0[113]. Spectra were searched against the human proteome, downloaded on 11.01.2021 for WCP and 21.05.2025 for PMP. Resulting peptide and protein tables were imported into Perseus[114] for downstream analysis. In Perseus, measured ion abundances were log_2_-transformed and data was filtered to remove common contaminants. PMP data was further filtered to only include proteins annotated with gene ontology cellular compartment (GOCC) terms plasma membrane (GO:0005886), cell surface (GO:0009986), or extracellular region (GO:0005576). WCP and PMP was further filtered to remove proteins not quantified in all three replicates of any one condition and missing values were imputed by fitting to a normal distribution. Replicates were grouped, and significantly altered proteins were identified using two-sample t-tests with an FDR threshold of <0.05. Fold-change and P-values were used to assign significance and to generate volcano plots, which were visualized in GraphPad Prism 9.

### Multi-electrode array (MEA)

Wells of a 24-well MEA CytoView plate (Axion BioSystems, M384-tMEA-24W) were coated with 500 μl 1× PLO and incubated overnight at room temperature. Wells were washed three times with sterile water and allowed to dry for ∼1 h before seeding 4 × 10^5^ 3 dpi neurons per well in 1 ml CN maintenance medium containing 1 μg/mL doxycycline. Cultures were maintained as described above with thrice weekly half-media changes. Recordings were performed on a Maestro Edge system (Axion BioSystems, 200-0888) using AxiS Navigator software. Activity and viability were routinely monitored with spikes detected using a threshold of 6× the standard deviation. On recording days, plates were equilibrated for >15 min at 37 °C and 5% CO_2_ before data acquisition. Spontaneous activity was recorded for 6 min in AxIS. Data were processed using NeuralMetric and the AxIS Plotting Tool before subsequent analysis in GraphPad Prism 9. Electrode bursts were identified using the ISI threshold algorithm in NeuralMetric with default parameters, minimum number of spikes = 5, max inter-spike interval = 100 ms. Network bursts were identified using the ISI threshold algorithm in NeuralMetric with default parameters, minimum number of spikes = 50, max inter-spike interval = 100 ms, minimum electrodes = 35%. Area under normalized cross-correlation (AUNCC) calculated using the cross-correlogram algorithm in NeuralMetric with default parameters, synchrony window = 20 ms.

## Author contributions

Conceptualization: H.G.B, Z.Z.H., J.E.D. Methodology: H.G.B, Z.Z.H, A.S.N., R.A, D.A.P. Investigation: H.G.B, Z.Z.H, A.S.N, S.S, D.A.N, J.O.S, R.A, D.tV. Visualization: H.G.B, Z.Z.H, J.E.D. Supervision: S.C.G, F.M.P, J.E.D. Writing—original draft: H.G.B, Z.Z.H, J.E.D. Writing—review & editing: H.G.B, Z.Z.H, J.O.S., D.tV., S.C.G., F.M.P., J.E.D.

## Data availability

The mass spectrometry proteomics data have been deposited to the ProteomeXchange Consortium via the PRIDE[115] partner repository with the dataset identifier PXD076397 (https://www.ebi.ac.uk/pride/archive/projects/PXD076397).

