## Supplementary Information for "Sialylated Gangliosides are Required for Plasma Membrane Organization and Neuronal Function"

**TITLE**

**SHORT TITLE**

Sialylated Gangliosides Organize Neuronal Membranes

**AUTHORS**

Henry G. Barrow*[,1,2], Zhuang Z. Han*[,1,2], Alex S. Nicholson[1,2], Sinéad Strasser[1], Daniel A. Nash[3], John O. Suberu[1], Robin Antrobus[1], Danielle te Vruchte[4], David A. Priestman[4], Stephen C. Graham[3], Frances M. Platt[4] and Janet E. Deane[1,2,^]

**This file contains:**

Supplementary Figures S1-S11

Supplementary Tables S1-S9

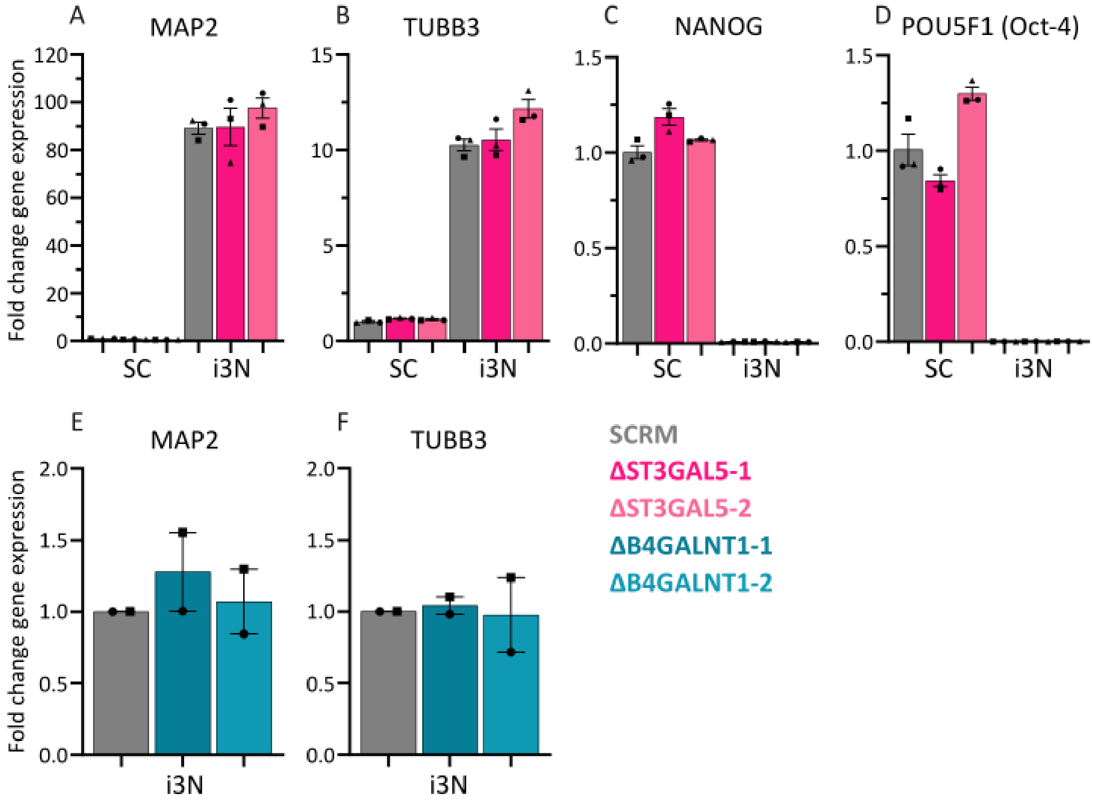


**Supplementary Figure S1.** A-D) qPCR analysis of gene expression for neuronal markers (MAP2, TUBB3 encoding β3-tubulin) and stem cell markers (NANOG, POU5F1 [OCT-4]) in SCRM and ΔST3GAL5 at 0 dpi (SC, stem cell) and 14 dpi (i3N). Fold change is calculated relative to 0 dpi SCRM controls, N=3 biological replicates (circles, squares, triangles) were carried out in technical triplicate. Mean ± SEM. E-F) qPCR analysis of gene expression for neuronal markers (MAP2, TUBB3) in SCRM and ΔB4GALNT1 cell lines at 28 dpi (i3N). Fold change is calculated relative to 28 dpi SCRM controls, N=2 biological replicates (circles, squares) were carried out in technical triplicate. Mean ± SEM.

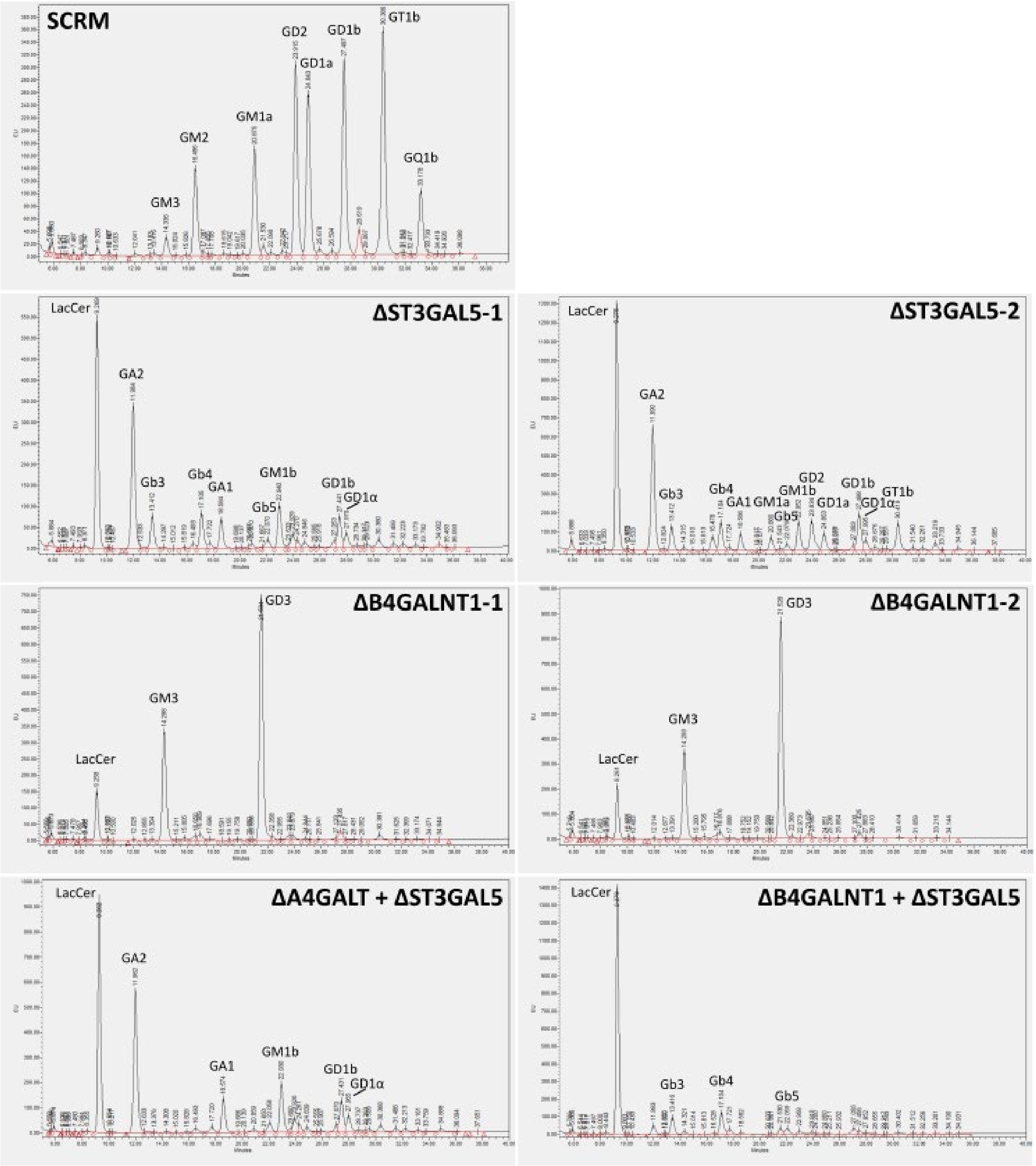


**Supplementary Figure S2.** Representative raw high-performance liquid chromatography (HPLC) traces for each cell line used in this study.

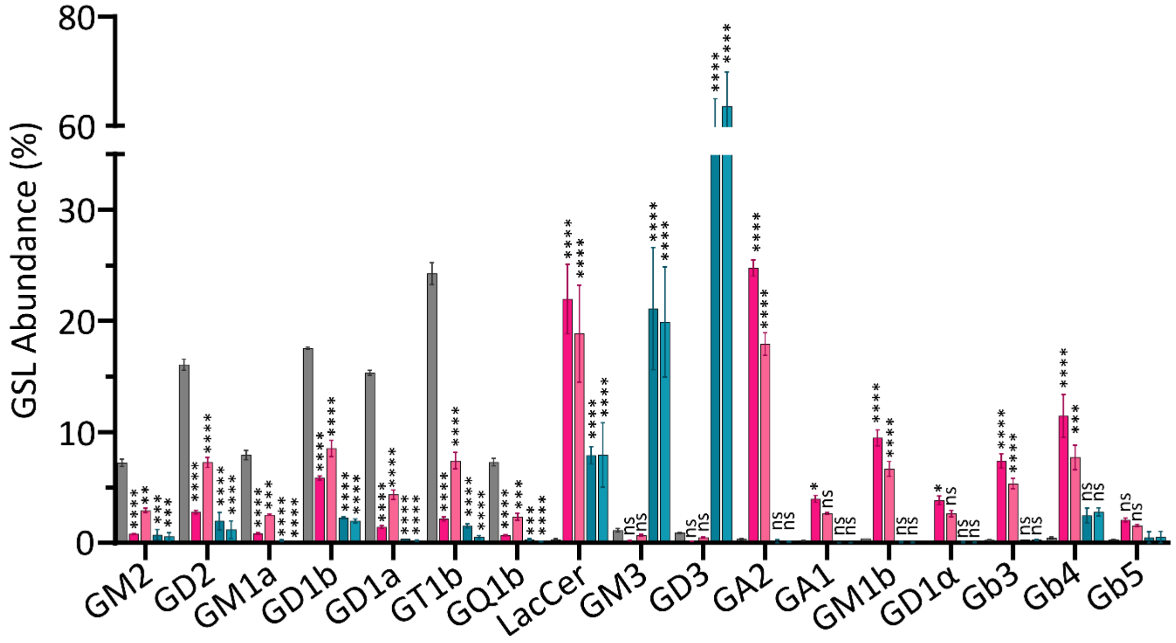


**Supplementary Figure S3.** Reproduction of manuscript figure panel 1E with statistics. Significance was calculated using two-way ANOVA with Dunnett's multiple comparisons testing against SCRM, ****$p$≤0.0001, ***$p$≤0.001, *$p$≤0.05, ns not significant. Mean ± SEM.

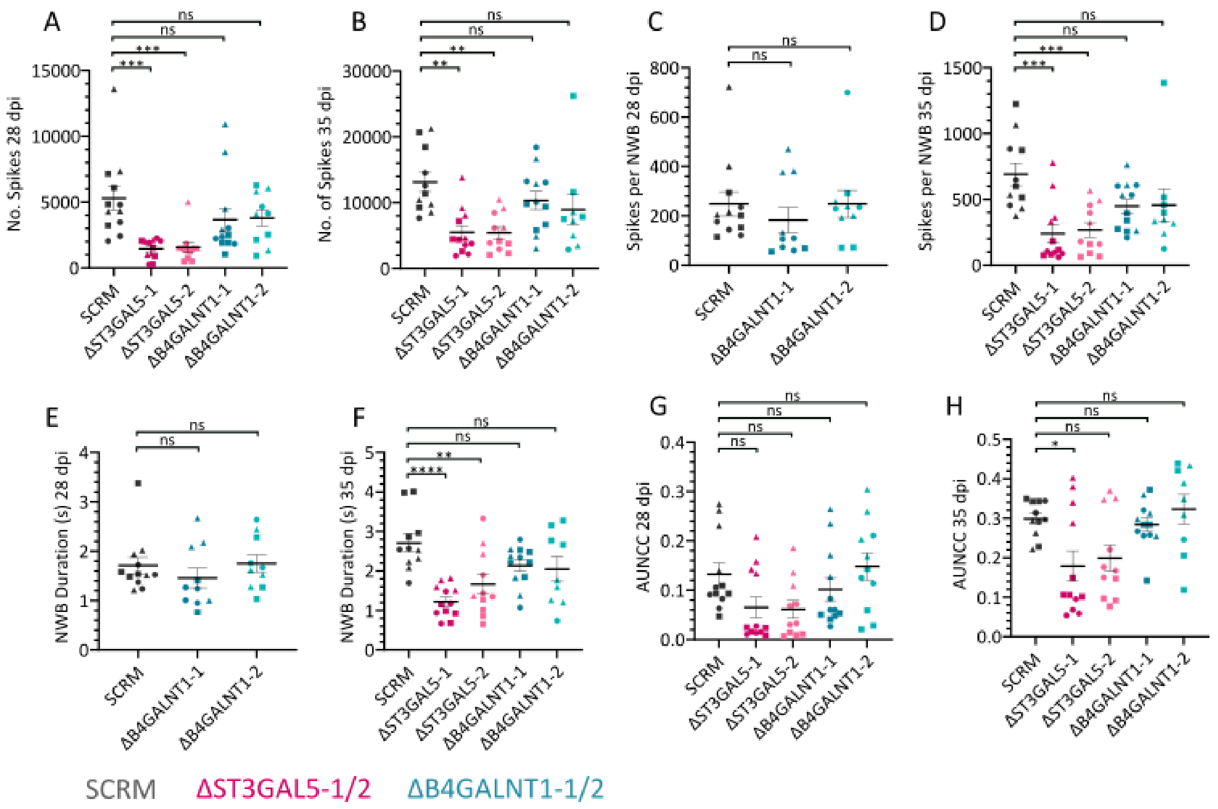


**Supplementary Figure S4.** Additional MEA analysis graphs: No. of spikes at 28 (A) or 35 (B) dpi, spikes per NWB at 28 (C) or 35 (D) dpi, NWB duration at 28 (E) or 35 (F) dpi, area under normalised cross-correlation (AUNCC) at 28 (G) or 35 (H) dpi. 9-12 wells measured per condition across N=3 biological replicates (circles, squares, triangles). Mean ± SEM. Significance calculated using one-way ANOVA with Dunnett's multiple comparisons testing against SCRM, ****$p \leq 0.0001$, ***$p \leq 0.001$, **$p \leq 0.01$, *$p \leq 0.05$, ns not significant.

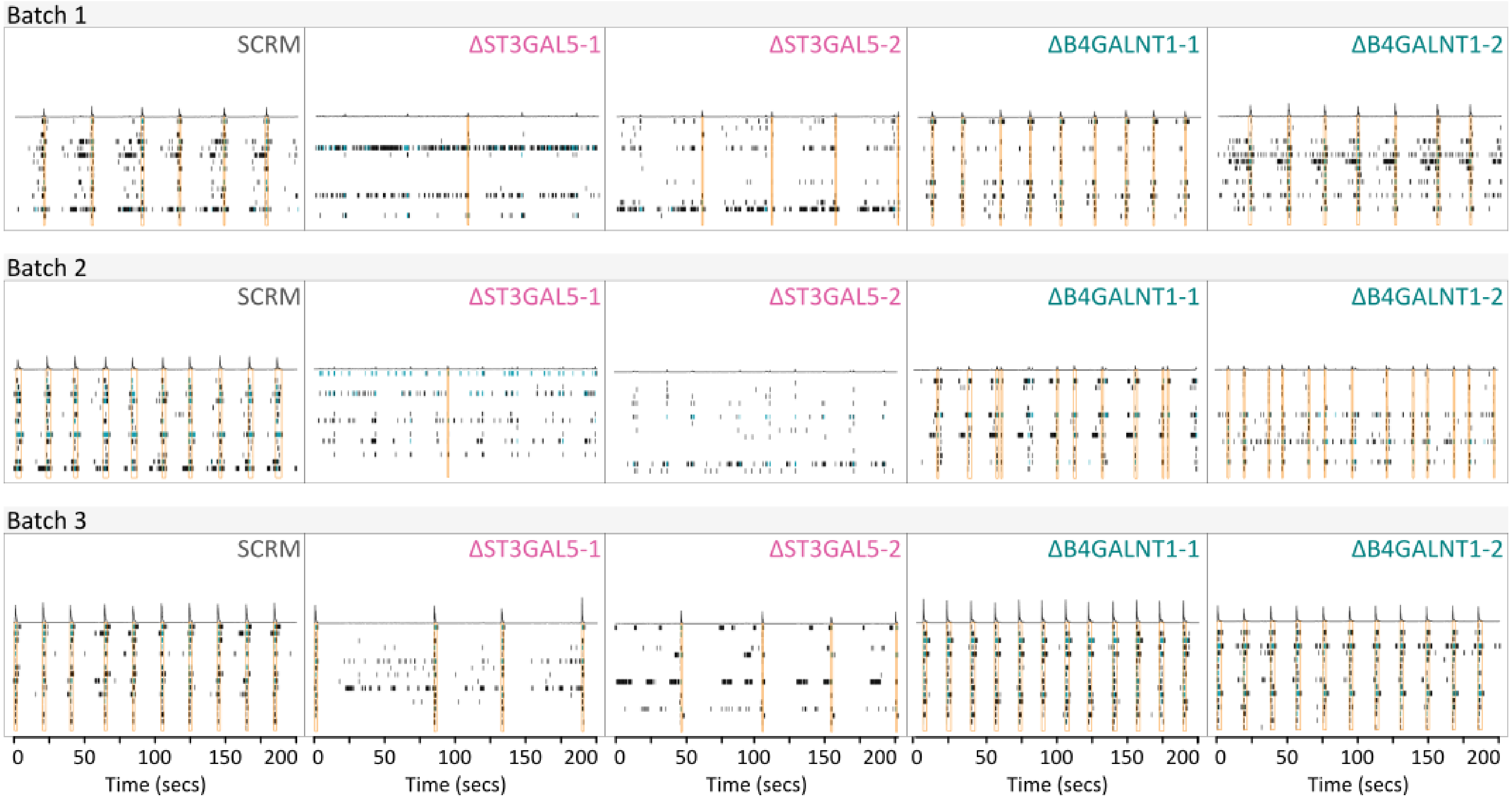


**Supplementary Figure S5.** Examples of raw MEA activity data (raster plots) from 28 dpi for all cell lines across N=3 independent experiments. Bursts (blue) and network bursts (orange) are highlighted.

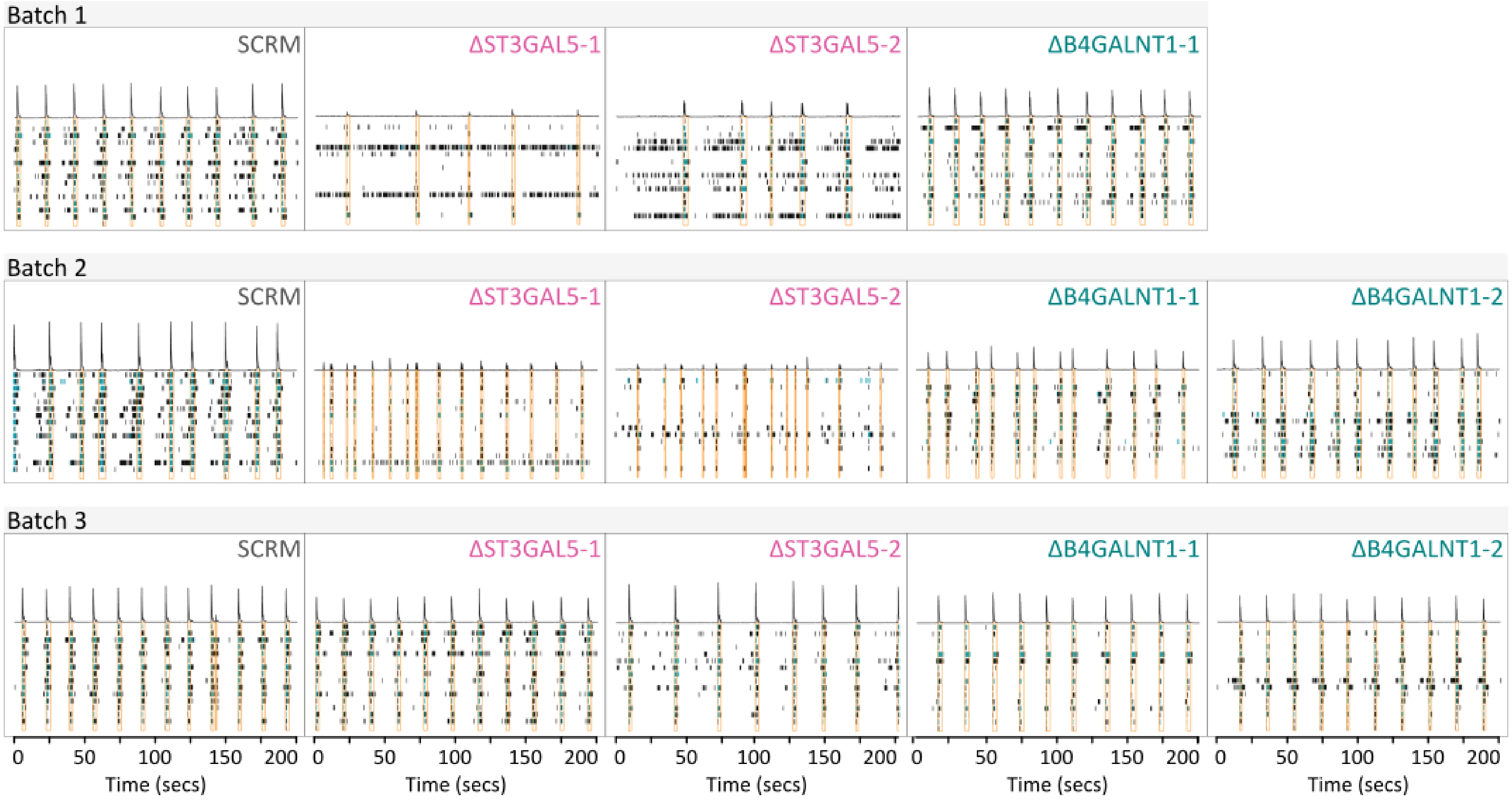


**Supplementary Figure S6.** Examples of raw MEA activity data (raster plots) from 35 dpi for all cell lines across N=2-3 independent experiments. Bursts (blue) and network bursts (orange) are highlighted.

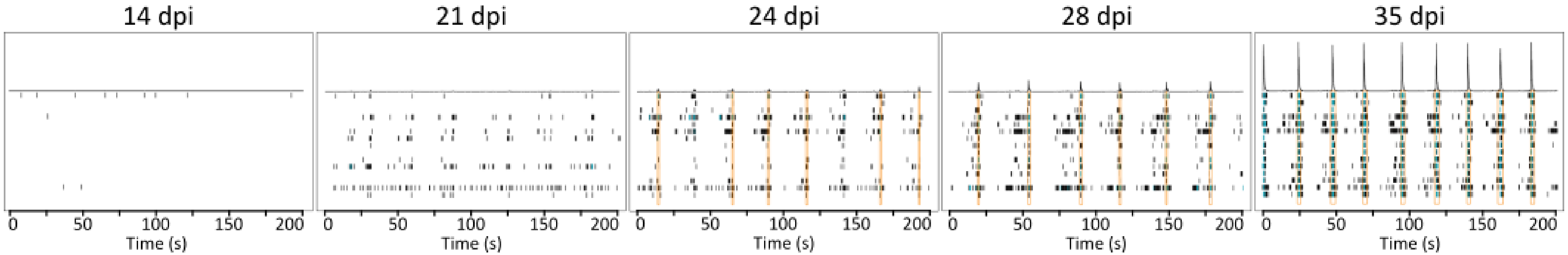


**Supplementary Figure S7.** Examples of raw MEA activity data (raster plots) over time demonstrating synchronised activity of SCRM cells by 28 dpi. Bursts (blue) and network bursts (orange) are highlighted.

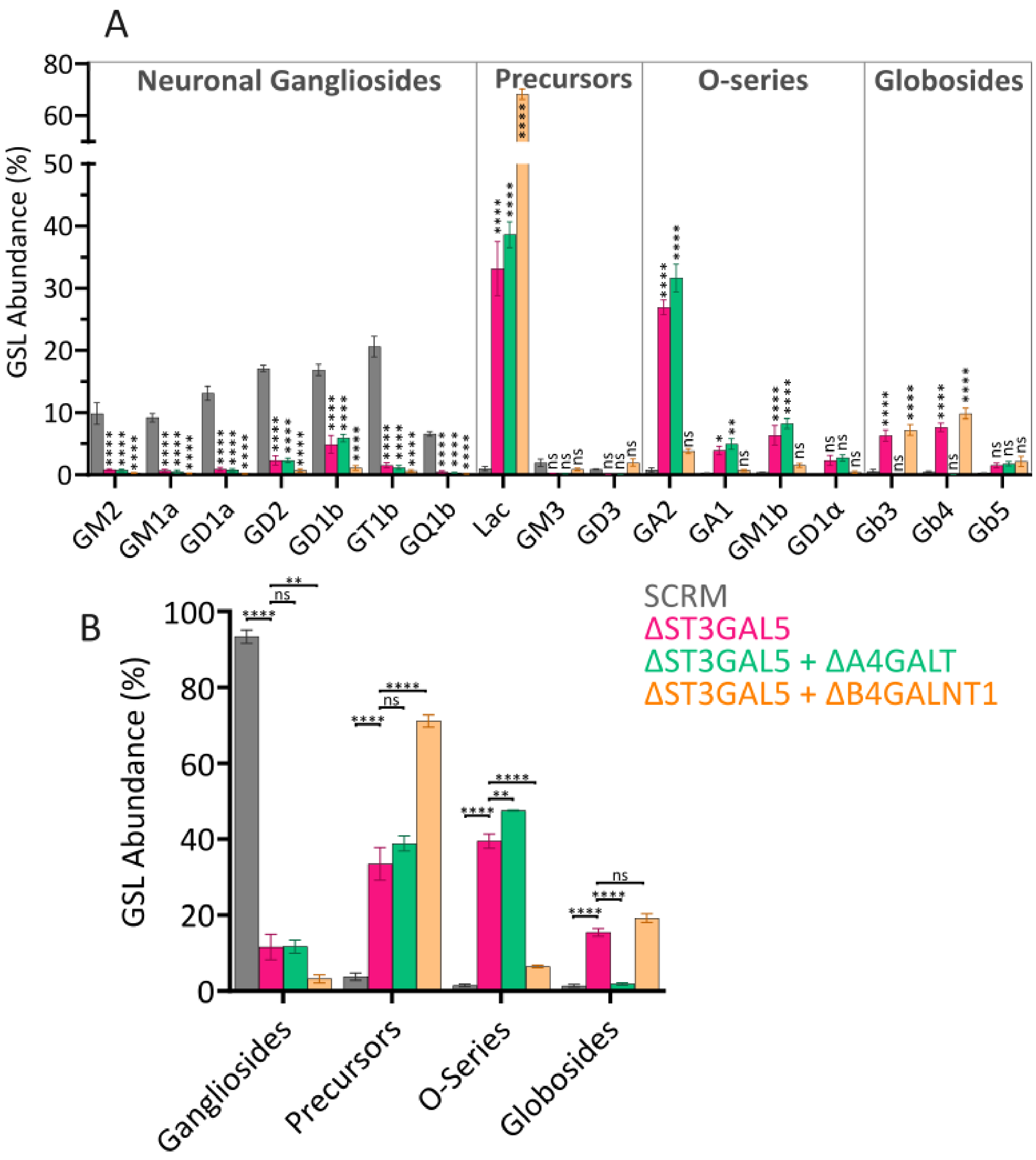


**Supplementary Figure S8.** A) Reproduction of manuscript figure panel 3C with statistics. B) Data from (A and Fig. 3C) grouped by lipid series into gangliosides (GM2, GD2, GM1a, GD1b, GD1a, GT1b, GQ1b), precursors (LacCer, GM3, GD3), o-series (GA2, GA1, GM1b, GD1α) and globosides (Gb3, Gb4, Gb5). Mean ± SEM. Significance was calculated using two-way ANOVA with Dunnett's multiple comparisons testing against SCRM, ****$p \leq 0.0001$, **$p \leq 0.01$, *$p \leq 0.05$, ns not significant.

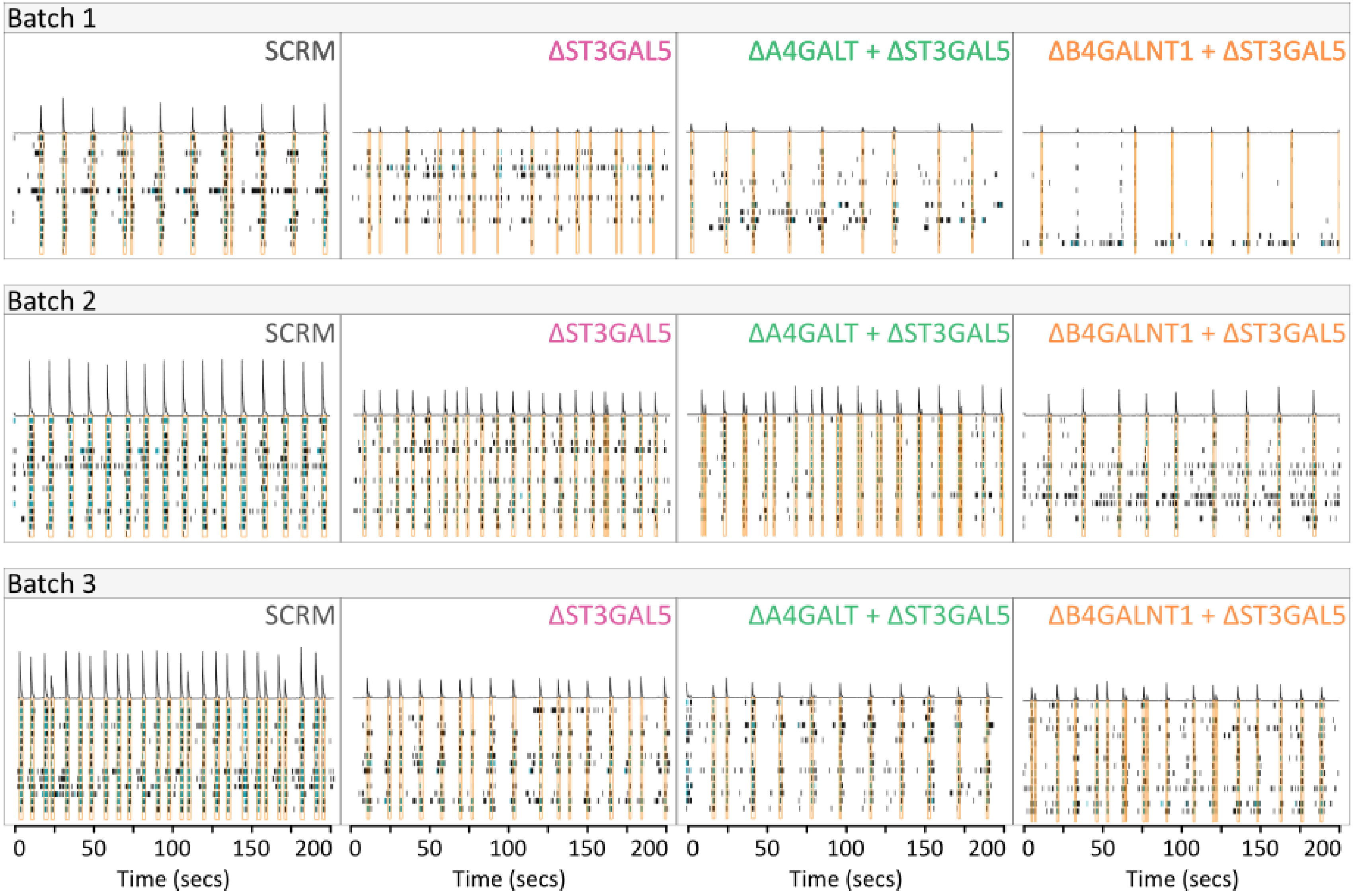


**Supplementary Figure S9.** Examples of raw MEA activity data (raster plots) from cell lines at 28 dpi across N=3 independent experiments.

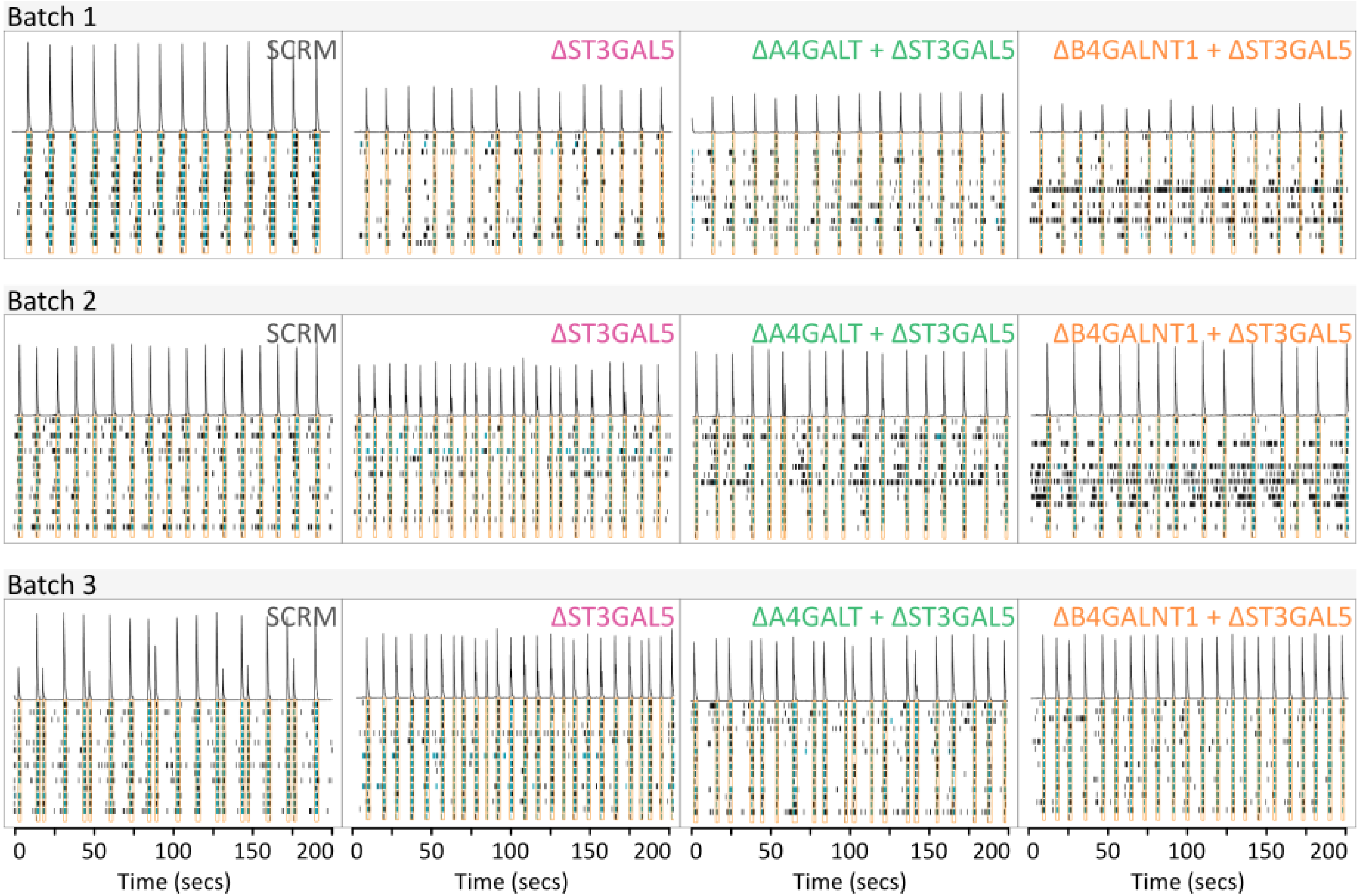


**Supplementary Figure S10.** Examples of raw MEA activity data (raster plots) from cell lines at 35 dpi across N=3 independent experiments.

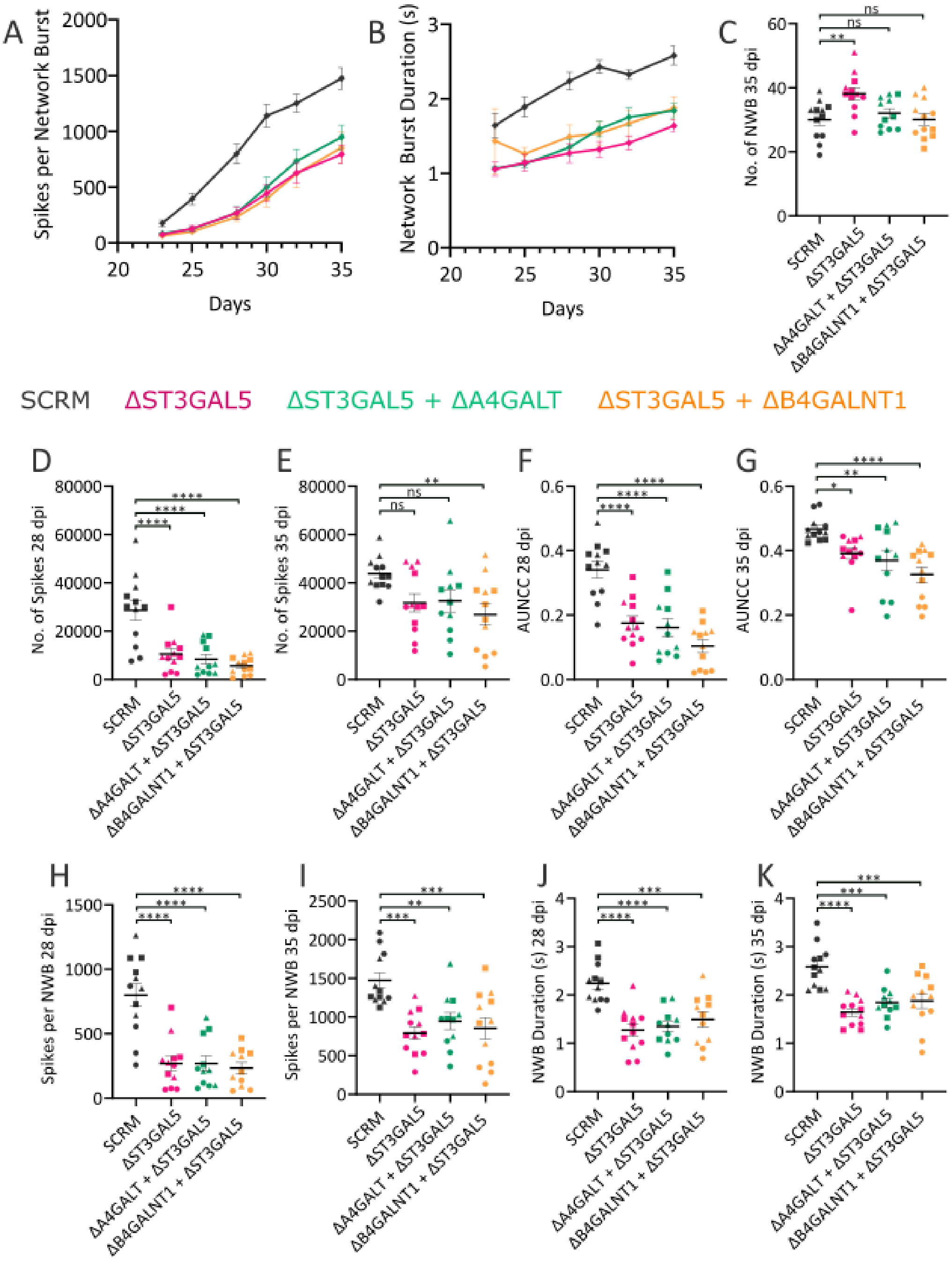


**Supplementary Figure S11. Additional MEA analysis graphs for ΔA4GALT+ΔST3GAL5 and ΔB4GALNT1+ΔST3GAL5 neurones.** A) Spikes per network burst and B) Network burst duration (s) over time. C) Number of network bursts at 35 dpi. Number of spikes at (D) 28 or (E) 35 dpi. Area under normalised cross correlation at (F) 28 or (G) 35 dpi. Spikes per network burst at (H) 28 or (I) 35 dpi. Network burst duration (s) at (J) 28 or (K) 35 dpi. Mean ± SEM are shown for 9-12 wells across N=3 biological replicates (circles, squares and triangles in C-K). Significance calculated using one-way ANOVA with Dunnett's multiple comparisons testing against SCRM, ****$p \leq 0.0001$, ***$p \leq 0.001$, *$p \leq 0.05$, ns not significant.

A

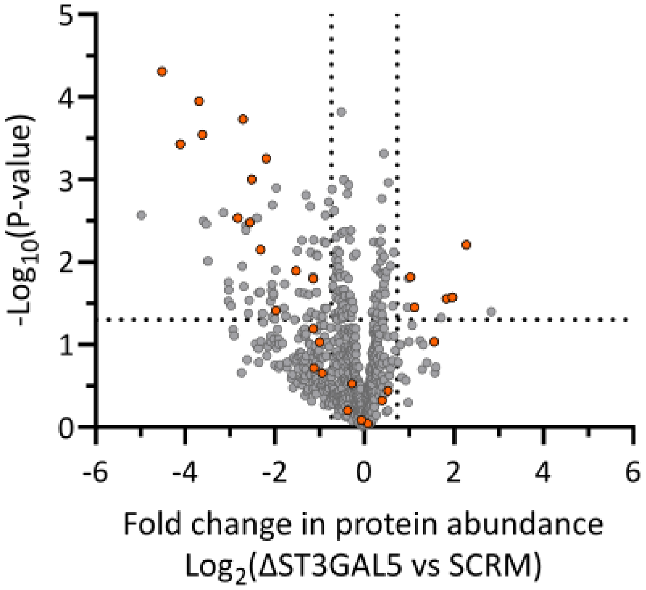


B

| Gene | Fold Change Decrease | P-value |
|---|---|---|
| NPY1R | 22.92 | 4.93E-05 |
| TMEM126A | 17.29 | 3.74E-04 |
| LIN7C | 12.92 | 1.13E-04 |
| RAB9B | 12.32 | 2.88E-04 |
| PHB2 | 7.11 | 2.93E-03 |
| CES5A | 6.55 | 1.88E-04 |
| HTR3A | 5.88 | 3.32E-03 |
| GALNT2 | 5.71 | 1.00E-03 |
| TOLLIP | 5.00 | 7.08E-03 |
| PFKL | 4.58 | 5.60E-04 |
| EXOC2 | 3.94 | 3.86E-02 |
| KCTD16 | 2.89 | 1.28E-02 |
| HCN2 | 2.21 | 1.60E-02 |
| COPB1 | 2.21 | 6.40E-02 |
| PALM | 2.18 | 1.92E-01 |
| ARL6IP5 | 2.00 | 9.27E-02 |
| PALM2AKAP2 | 1.93 | 2.21E-01 |
| ABCA3 | 1.29 | 6.32E-01 |
| PEG10 | 1.22 | 2.96E-01 |
| NBEA | 1.05 | 8.18E-01 |

C

| Gene | Fold Change Increase | P-value |
|---|---|---|
| DGAT1 | 4.83 | 6.20E-03 |
| DTNA | 3.89 | 2.68E-02 |
| NHERF2 | 3.55 | 2.78E-02 |
| ZYX | 2.95 | 9.20E-02 |
| SLC4A2 | 2.17 | 3.54E-02 |
| LY6H | 2.03 | 1.51E-02 |
| SYNGR1 | 1.44 | 3.62E-01 |
| BLTP1 | 1.31 | 4.75E-01 |
| BCR | 1.05 | 9.04E-01 |

**Supplementary Figure S12. Imputation in ΔST3GAL5 PMP samples.** A) Proteins highlighted in orange were quantified in all three SCRM but zero ΔST3GAL5 PMP samples. B/C) List of proteins that were quantified in all three SCRM but zero ΔST3GAL5 PMP samples that after imputation are considered to be (B) decreased or (C) increased in abundance in ΔST3GAL5 vs SCRM PMP samples. If a protein is quantified only in the SCRM samples then it is likely decreased in abundance in the ΔST3GAL5 samples and those considered here to be increased are likely an artefact of imputation. If a protein is quantified to very low abundance in SCRM but absent in ΔST3GAL5 then values may be imputed for ΔST3GAL5 above those of SCRM.

### SUPPLEMENTARY TABLES

**Table S1**. Guide RNA target sequences.

| sgRNA | Sequence |
|---|---|
| ΔST3GAL5-1 | GCGGCGGCCCCCAGCTGAAT |
| ΔST3GAL5-2 | GGCGCCCCCTCATTAGTATG |
| ΔB4GALNT1-1 | GCTCAGCTCCCGGCTCGTTG |
| ΔB4GALNT1-2 | CGCGTCCCACTGGCAGCGCG |
| ΔA4GALT-1 | CCGCTGGAGCTAGAGGTACG |
| ΔA4GALT-2 | GCTGGAGCTAGAGGTACGAG |

**Table S2.** PM proteins decreased in abundance in ΔST3GAL5 neurons. G-protein coupled receptors (orange), ion channels (green) and synaptic adhesion proteins (pink) are highlighted.

| Gene | Fold Change Decrease | P-value |
|---|---|---|
| KISS1R | 31.56 | 2.71E-03 |
| NPY1R | 22.92 | 4.93E-05 |
| TMEM126A | 17.29 | 3.74E-04 |
| LIN7C | 12.92 | 1.13E-04 |
| RAB9B | 12.32 | 2.88E-04 |
| SLC2A13 | 12.07 | 3.19E-03 |
| EXOC6B | 11.58 | 3.44E-03 |
| EXOC7 | 11.28 | 9.69E-03 |
| CXCR4 | 8.88 | 2.52E-03 |
| ENO1 | 8.17 | 1.75E-02 |
| PTPRN | 8.17 | 2.98E-02 |
| DCD | 8.14 | 2.19E-02 |
| NRXN3 | 7.88 | 3.39E-02 |
| PHB2 | 7.11 | 2.93E-03 |
| EFR3B | 6.64 | 1.12E-02 |
| CES5A | 6.55 | 1.88E-04 |
| SLC26A6 | 6.48 | 1.97E-02 |
| ATP11A | 6.26 | 3.62E-03 |
| PLPP2 | 6.26 | 4.13E-03 |
| CDK5 | 6.01 | 4.19E-02 |
| TRHR | 5.93 | 2.93E-02 |
| HTR3A | 5.88 | 3.32E-03 |
| GALNT2 | 5.71 | 1.00E-03 |
| NF1 | 5.29 | 2.94E-03 |
| SLC44A3 | 5.12 | 2.35E-02 |

| | | |
|---|---|---|
| KCNH7 | 5.03 | 2.32E-02 |
| TOLLIP | 5.00 | 7.08E-03 |
| PKM | 4.83 | 4.19E-02 |
| PFKL | 4.58 | 5.60E-04 |
| EXOC1 | 4.27 | 3.01E-02 |
| VDAC1 | 4.20 | 2.20E-02 |
| PPM1A | 4.19 | 1.86E-02 |
| SEZ6 | 4.19 | 4.66E-02 |
| PTGER3 | 4.16 | 2.04E-03 |
| MYADM | 4.12 | 2.58E-02 |
| LGI1 | 4.08 | 2.42E-02 |
| NPDC1 | 4.01 | 4.88E-02 |
| CLCN4 | 3.96 | 1.24E-02 |
| EXOC2 | 3.94 | 3.86E-02 |
| OPRL1 | 3.92 | 1.26E-03 |
| SCAMP1 | 3.47 | 3.61E-02 |
| EXOC8 | 3.44 | 1.85E-02 |
| KIAA1549 | 3.38 | 4.55E-02 |
| AMER2 | 2.95 | 5.00E-02 |
| KCTD16 | 2.89 | 1.28E-02 |
| NPR3 | 2.83 | 7.33E-03 |
| EXOC4 | 2.74 | 2.34E-02 |
| EPHA2 | 2.64 | 5.48E-03 |
| PRDX5 | 2.53 | 2.40E-02 |
| LRRTM3 | 2.48 | 1.55E-03 |
| TPI1 | 2.44 | 4.69E-02 |
| CLASP2 | 2.37 | 2.50E-02 |
| CAND1 | 2.32 | 2.13E-03 |
| FIGNL1;VPS4A;VPS4B | 2.30 | 7.62E-03 |
| CKAP5 | 2.30 | 4.04E-02 |
| SLC22A5 | 2.22 | 1.50E-02 |
| HCN2 | 2.21 | 1.60E-02 |
| MTOR | 2.19 | 5.41E-03 |
| CHRM4 | 2.11 | 4.36E-02 |
| KCNA3 | 2.10 | 2.32E-02 |
| ENDOD1 | 2.09 | 8.17E-03 |
| SLC7A8 | 2.08 | 8.44E-03 |
| SERINC3 | 1.94 | 5.00E-02 |
| RNH1 | 1.92 | 4.56E-02 |
| SLCO3A1 | 1.88 | 5.69E-03 |
| FRRS1L | 1.86 | 3.67E-02 |
| RTN4RL1 | 1.84 | 2.70E-03 |
| ATP6V1H | 1.82 | 5.91E-03 |
| EPRS1 | 1.81 | 3.66E-02 |
| TPBG | 1.74 | 1.87E-03 |

| | | |
|---|---|---|
| NPEPPS | 1.66 | 2.21E-02 |

**Table S3.** PM proteins increased in abundance in ΔST3GAL5 neurons.

| Genes | Fold Change Increase | P-value |
|---|---|---|
| FADS2 | 7.12 | 4.00E-02 |
| DGAT1 | 4.83 | 6.20E-03 |
| DTNA | 3.89 | 2.68E-02 |
| NHERF2 | 3.55 | 2.78E-02 |
| CA11 | 3.27 | 4.69E-02 |
| SLC4A2 | 2.17 | 3.54E-02 |
| LY6H | 2.03 | 1.51E-02 |
| USP12;USP46 | 1.98 | 1.58E-02 |
| APOC3 | 1.95 | 3.54E-02 |

**Table S4.** PM proteins decreased in abundance in ΔB4GALNT1 neurons.

| Gene | Fold Change Decrease | (P-value) |
|---|---|---|
| MTOR | 6.06 | 5.00E-02 |
| PHB1 | 2.34 | 3.19E-02 |
| EXOC6B | 1.84 | 6.85E-03 |
| CAND1 | 1.69 | 1.64E-03 |
| EXOC7 | 1.67 | 1.26E-03 |

**Table S5.** PM proteins increased in abundance in ΔB4GALNT1 neurons.

| Genes | Fold Change | P-value |
|---|---|---|
| H2BC11;H2BCx | 2.28 | 4.89E-02 |
| LGI2 | 1.70 | 1.19E-02 |

**Table S6.** Proteins increased in abundance at whole cell level in ΔST3GAL5 neurons.

| Gene | Fold Change | P-value |
|---|---|---|
| SCGN | 1.88 | 9.06E-04 |
| PDLIM1 | 1.72 | 1.89E-02 |

**Table S7.** Proteins decreased in abundance at whole cell level in ΔST3GAL5 neurons.

| Gene | Fold Change | P-value |
|---|---|---|
| ANR12 | -1.76 | 4.98E-02 |

**Table S8.** Proteins increased in abundance at whole cell level in ΔB4GALNT1neurons.

| Gene | Fold Change | P-value |
|---|---|---|
| MT-CO2 | 1.84 | 3.62E-02 |
| POTEF | 1.80 | 7.59E-03 |
| CD81 | 1.69 | 4.88E-02 |

**Table S9**: Taqman gene expression assay probe sets.

| Gene | Probe |
|---|---|
| GAPDH | Hs02786624_g1 |
| ST3GAL5 | Hs01105377_m1 |
| B4GALNT1 | Hs011100791_g1 |
| A4GALT | Hs05058505_s1 |
| NANOG | Hs02387400_g1 |
| OCT-4 | Hs04260367_gH |
| MAP2 | Hs00258900_m1 |
| TUBB3 | Hs00801390_s1 |
